# Tracheal tuft cell-released leukotrienes promote antibacterial immune responses

**DOI:** 10.1101/2025.10.08.681191

**Authors:** Mohamed Ibrahem Elhawy, Noran Abdel Wadood, Maria Grammer, Paula Gehrich, Emely Herrmann, Giuseppina Sole Lanzilli, Firat Biradli, Caroline Klozenbücher, Andreas Klein, Nosaibah Alakkam, Monika I. Hollenhorst, Saskia B. Evers, Soumya Kusumakshi, Amanda Wyatt, Andreas M. Kany, Sören L. Becker, Markus Bischoff, Veit Flockerzi, Thomas Gudermann, Vladimir Chubanov, Martin Empting, Anna K. H. Hirsch, Christoph Schneider, Ulrich Boehm, Gabriela Krasteva-Christ

## Abstract

Tuft cells act as crucial sentinels in the airways that detect bacterial metabolites. In response, tuft cells release signaling molecules that trigger immune responses essential for clearing the infection. The molecular mechanisms driving immune cell activation following tuft cell stimulation in pneumonia are still not fully understood. Here, we identify tuft cells as the primary source of proinflammatory leukotrienes (LTs), which are released in the presence of pathogenic bacteria in the airways. We show that tracheal tuft cells discriminate pathogenic from non-pathogenic bacteria by sensing adenosine triphosphate (ATP) released from pathogens such as *Pseudomonas aeruginosa* and *Rodentibacter pneumotropicus* within the first 4 h of invasion, and recruit neutrophils and macrophages to the trachea and alveolar spaces. Taste signaling through the chemosensory transient receptor potential cation channel subfamily M member 5 (Trpm5) channel was essential for tuft cell activation and LT release. Mice lacking Trpm5 were not capable of detecting bacteria-released ATP and became colonized upon *R. pneumotropicus* infection. In contrast, Trpm5+/+ mice cleared the pathogen. We uncover a critical tuft cell-dependent sensing mechanism in pneumonia and establish tracheal tuft cells as both detectors of bacterial extracellular ATP and triggers of acute innate immune responses.

## INTRODUCTION

Bacterial respiratory tract infections represent a global public health concern. Although pathogen detection by respiratory epithelial cells and the subsequent activation of immune cells are essential for resolving these infections, the underlying mechanisms driving the protective responses remain largely unclear. Recent studies have identified tuft cells as key regulators of innate immunity in the airways during *Pseudomonas aeruginosa* infection^1–3^. Tuft cells are specialized epithelial cells located at mucosal surfaces throughout the body^4^. In the trachea, tuft cells act as sensors for bacterial signaling molecules, including *P. aeruginosa* quorum-sensing molecules (QSM) and formyl peptides^2,5^ activating the canonical taste signal transduction cascade, particularly the calcium-activated monovalent-specific ion channel Trpm5^6–9^. Tracheal tuft cells also detect metabolites such as succinate^10^, which increases in the bronchoalveolar lavage fluid (BALF) during bacterial infection^11^. In response, tracheal tuft cells release acetylcholine (ACh) in a Trpm5-dependent manner^5,8,10^. The tuft cell-derived ACh exerts paracrine effects on adjacent epithelial cells and sensory nerve fibers, thereby enhancing mucociliary clearance^5,7,8,10,12^ and triggering neutrophil recruitment^2^ and protective respiratory reflexes^13^.

Recently, we have shown that tracheal tuft cell activation also induces a Trpm5-dependent release of ATP, followed by dendritic cell (DC) recruitment and activation^3^. Although tuft cell-released ATP contributes to host defense and is required to combat *P. aeruginosa* infection^3^, its specific role in initiating acute immune responses remains poorly understood. It appears plausible that ATP derived from tuft cells acts in an autocrine manner to induce or sustain protective immune responses. This hypothesis is supported by the expression of purinergic receptors in tracheal tuft cells^14–16^. Solitary chemosensory cells (SCCs) of the nose share a common transcription profile with tracheal tuft cells and respond to extracellular ATP (eATP) or aeroallergens by releasing cysteinyl leukotrienes (CysLTs)^17^. In *Alternaria alternata* infection, CysLTs are released by tuft cells and contribute to airway type 2 inflammation by inducing eosinophilia in the alveolar spaces^17^. Tuft cell-derived CysLTs also play a crucial role in regulating immune responses in the gut in response to infection with the intestinal parasites *Heligmosomoides polygyrus* or *Nippostrongylus brasiliensis*^18^. However, it is still unknown whether ATP released by tuft cells in response to bacterial metabolites or bacterial ATP induces a leukotriene (LT) release in the trachea, and most importantly, whether this is crucial to mount immune responses in the airways in response to bacterial infection.

LTs are lipid metabolites of arachidonic acid (AA). Initially, they were considered to be generated primarily by immune cells, including macrophages, neutrophils, and mast cells^19–21^. However, several reports suggested that LTs can also be produced by epithelial cells^22,23^. LT synthesis is based on a common enzymatic pathway leading to the generation of two main categories: leukotriene B4 (LTB_4_) and CysLTs, which include leukotriene C_4_ (LTC_4_), D_4_ (LTD_4_), and E_4_ (LTE_4_)^24^. Upon cellular activation, arachidonic acid (AA) is released from cellular membranes in a reaction mediated by the enzyme group IV cytosolic phospholipase A2. The released AA is then oxidized by arachidonate 5-lipoxygenase (ALOX5) in the presence of the ALOX5-activating protein (ALOX5-AP; also known as FLAP) to generate LTA_4_, which is subsequently converted by the LTA4 hydrolase (LTA4H) into LTB_4_ or by leukotriene C_4_ synthase (LTC4S) into LTC_4_^25^. Both LTB4 and LTC4 are transported into the extracellular milieu, where LTC_4_ is converted into the two final metabolites LTD_4_ and LTE_4_^26,27^. Both LTB_4_ and CysLTs have been reported to play a role in various immunological processes. LTB_4_ exerted a chemotactic effect on neutrophils in the lungs of mice infected with *Klebsiella pneumoniae*^28^. CysLTs promoted nasal clearance of *Streptococcus pneumoniae* in infected allergic mice^29^. While these findings implicate LTs in modulating innate immune responses during bacterial infections, it remains unknown whether they contribute to tuft cell-mediated immunity.

Here, we investigated how tuft cell-derived LTs contribute to innate immune responses combating bacterial infection. We further tested whether tracheal tuft cells act as sensors for bacterial-derived extracellular ATP (eATP) released during the early stages of bacterial infection and defined the interplay between eATP sensing and LT release by tracheal tuft cells. Finally, we investigated whether the eATP-sensing capacity of tracheal tuft cells enables them to discriminate between commensal and pathogenic bacteria in the airways.

## RESULTS

### Tracheal tuft cells release CysLTs and LTB_4_

During bacterial infection, tracheal tuft cells initiate rapid and tailored immune responses by releasing signaling molecules, e.g., ACh and ATP^2,3,5^. Nasal SCCs generate CysLTs in response to aeroallergens^17^, but it remains unclear whether tracheal tuft cells also engage LT release to mediate protective immune responses during bacterial infection. To examine the contribution of tracheal tuft cells to LT production, including both CysLTs and LTB_4_, we performed qRT-PCR on tracheal tissue collected from wild-type mice (*Trpm5^+/+^*) and *Trpm5*-DTA mice. In this mouse strain, Trpm5^+^ cells are genetically ablated via diphtheria toxin A (DTA)^2,30^. Immunohistochemistry for Trpm5 and qRT-PCR analysis in tracheae from *Trpm5*-DTA mice and in *Chat*-eGFP mice, in which tracheal tuft cells can be identified by their green fluorescence^3^, confirmed the complete absence of Trpm5^+^ tuft cells and *Trpm5* expression in the *Trpm5*-DTA mouse model (Figures S1A-C), indicating the specific expression of Trpm5 only in tuft cells in the trachea. Our qRT-PCR analyses revealed that the mRNA levels of LT-synthesizing genes *Alox5*, *Alox5ap*, and *Ltc4s* were significantly decreased in tracheae of mice lacking tuft cells compared to wild-type (*Trpm5^+/+^*) mice, whereas the relative expression of *Pla2gra* and *Lta4h* remained unchanged (Figures S1D-H). We confirmed our findings through *in silico* analysis of published single-cell RNA sequencing data (dataset GSE116525)^16^ (Figures S2A-E) and further validated them using qRT-PCR of FACS-sorted primary tracheal tuft cells from *Chat-*eGFP mice (Figures S2F-J). The specificity of the FACS-sorting strategy was validated by detecting *Trpm5*-mRNA exclusively in tracheal tuft cells (EpCAM^+^, GFP^+^) and not in other tracheal epithelial cell types (EpCAM^+^, GFP^-^) or immune cells (EpCAM^-^, CD45^+^) (Figures S3A and B). Next, we asked whether specific stimulation of tracheal tuft cells, involving a rise in intracellular Ca²⁺ and thus mimicking an activation of tuft cells through stimulation of type 2 taste receptors (Tas2R), induces a release of LTB_4_ and CysLTs. We used a *Trpm5*-DREADD;tGFP mouse model^2,30^, in which tuft cells express designer receptors exclusively activated by designer drugs (DREADD) (Figure 1A). Following treatment of *Trpm5*-DREADD tracheae with the synthetic DREADD ligand clozapine N-oxide (CNO) in a final concentration of 100 µM, we detected significantly increased levels of CysLTs and LTB_4_ in tracheal supernatants (Figures 1B and C). Our findings indicate that tracheal tuft cell activation leads to the release of CysLTs and LTB_4_, consistent with the exclusive expression of *Alox5* and *Alox5ap* in tuft cells (Figure 1D).

**Fig. 1:**
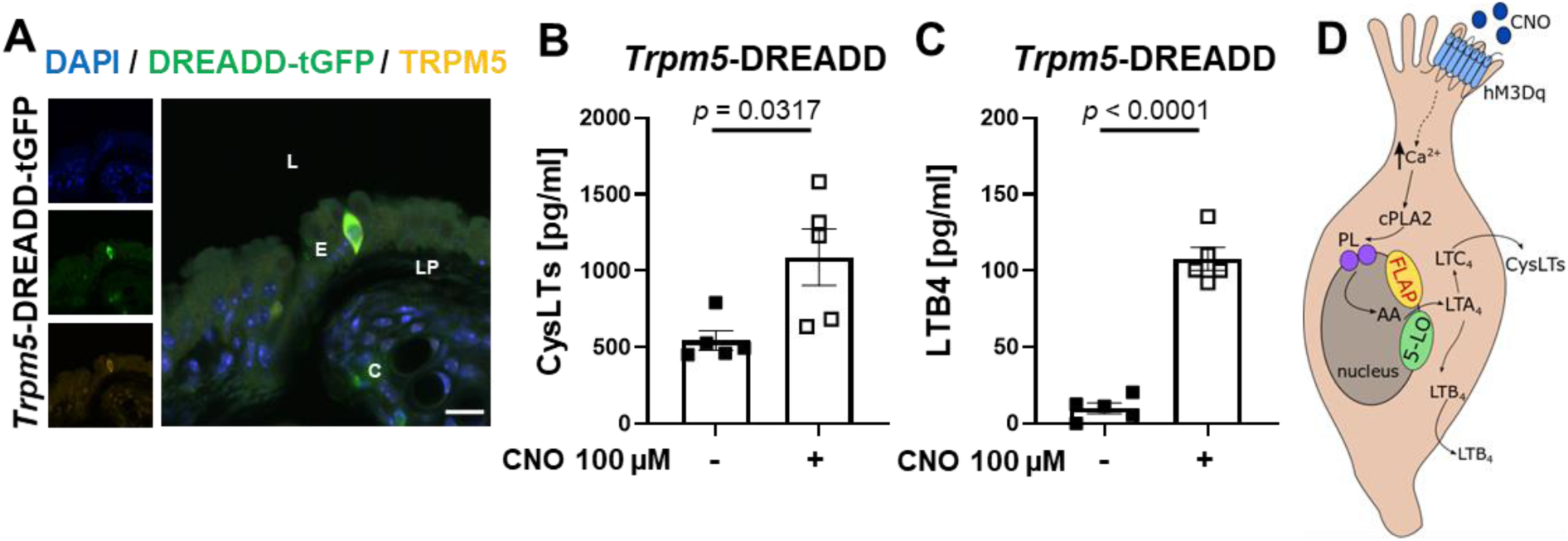
Tracheal tuft cells release CysLTs and LTB4 upon stimulation. (A) Representative immunohistochemical staining of a mouse *Trpm5*-DREADD-tGFP trachea. Scale bar, 20 µm. L, lumen; E, epithelium; LP, lamina propria; C, cartilage. (B-C) ELISA measurements of CysLTs and LTB4 in tracheal supernatants collected from *Trpm5*-DREADD-tGFP mice (n=5) stimulated either with 100 µM CNO or with vehicle. (D) Schematic drawing of the release of LTs from tracheal tuft cells. hM3Dq; Gq-coupled human M3 muscarinic receptor, CysLTs; cysteinyl leukotrienes. Data are shown as mean ± SEM of 5 independent experiments. Two-tailed unpaired Student’s *t*-test (B-C).

### Tracheal tuft cells’ LT release depends on Trpm5 and ATP

Tracheal tuft cells express Tas2R along with the downstream taste transduction cascade^8,14,15^, enabling them to detect luminal stimuli. Tuft cell activation with Tas2R agonists, including denatonium and N-acyl homoserine lactones (AHLs) from *P. aeruginosa*^31^, induces a release of ACh and ATP, and evokes protective immune responses^2,3,5,8^. These effects were prominent in tracheae of *Trpm5^+/+^* but not of *Trpm5^-/-^*mice^2,3^, indicating that Trpm5 signaling in tracheal tuft cells plays a central role in regulating tuft cell-mediated immune responses. To determine whether Trpm5 signaling in tuft cells contributes to LT release, we measured CysLTs and LTB_4_ concentration upon stimulation with Tas2R agonists. Treatment of freshly isolated tracheae with 1 mM denatonium triggered CysLT and LTB_4_ release in the *Trpm5^+/+^*, but not in *Trpm5^-/-^* mice (Figures 2A and B). This denatonium concentration was previously found to be specific for tracheal tuft cell activation^2,8^. Using the *Trpm5*-DTA model, we further demonstrated that tuft cells are mandatory for LT generation in the tracheal epithelium, as denatonium (1 mM) failed to trigger CysLT or LTB_4_ release in the absence of tuft cells (Figures S4A and B). Moreover, the FLAP inhibitor MK-886 (1 µM) reduced the elevated CysLT and LTB_4_ levels induced by denatonium in *Trpm5^+/+^* mice to those detected in the non-stimulated controls (Figures 2A and B). Notably, CysLT and LTB4 baseline levels did not differ between tracheae from *Trpm5^+/+^* and *Trpm5^-/-^* mice. Similarly, incubation of tracheae with 50 µM *Pseudomonas* quinolone signal (PQS), a concentration previously shown to specifically activate tracheal tuft cells^2^, induced CysLT and LTB_4_ release, which was significantly reduced after treatment with 1 µM MK-886, restoring levels to baseline (Figures 2C and D). Together, these results indicate that tracheal tuft cells release CysLTs and LTB_4_ upon stimulation with denatonium or bacterial metabolites. Based on our previous observation that tracheal tuft cells release ATP in response to stimulation with 1 mM deantonium^3^, and the fact that tracheal tuft cells express purinergic receptors^14–16^, we hypothesized that ATP may act in an autocrine manner to induce CysLT and LTB_4_ release. Treatment of tracheae with the ATP-hydrolyzing enzyme apyrase (5 U/ml) abolished CysLT and LTB_4_ release in response to stimulation with denatonium (Figures 2E and F), confirming that tuft cell-derived ATP indeed promotes their release. Further supporting this, stimulation of explanted tracheae with 1 mM ATP resulted in a robust CysLT and LTB_4_ release (Figures 2E and F). This concentration of ATP was previously shown to induce a release of CysLTs from nasal SCCs^17^. Together, these findings indicate that the tuft cell-released ATP acts in an autocrine manner to trigger the release of CysLTs and LTB_4_ (Figure 2G).

**Fig. 2:**
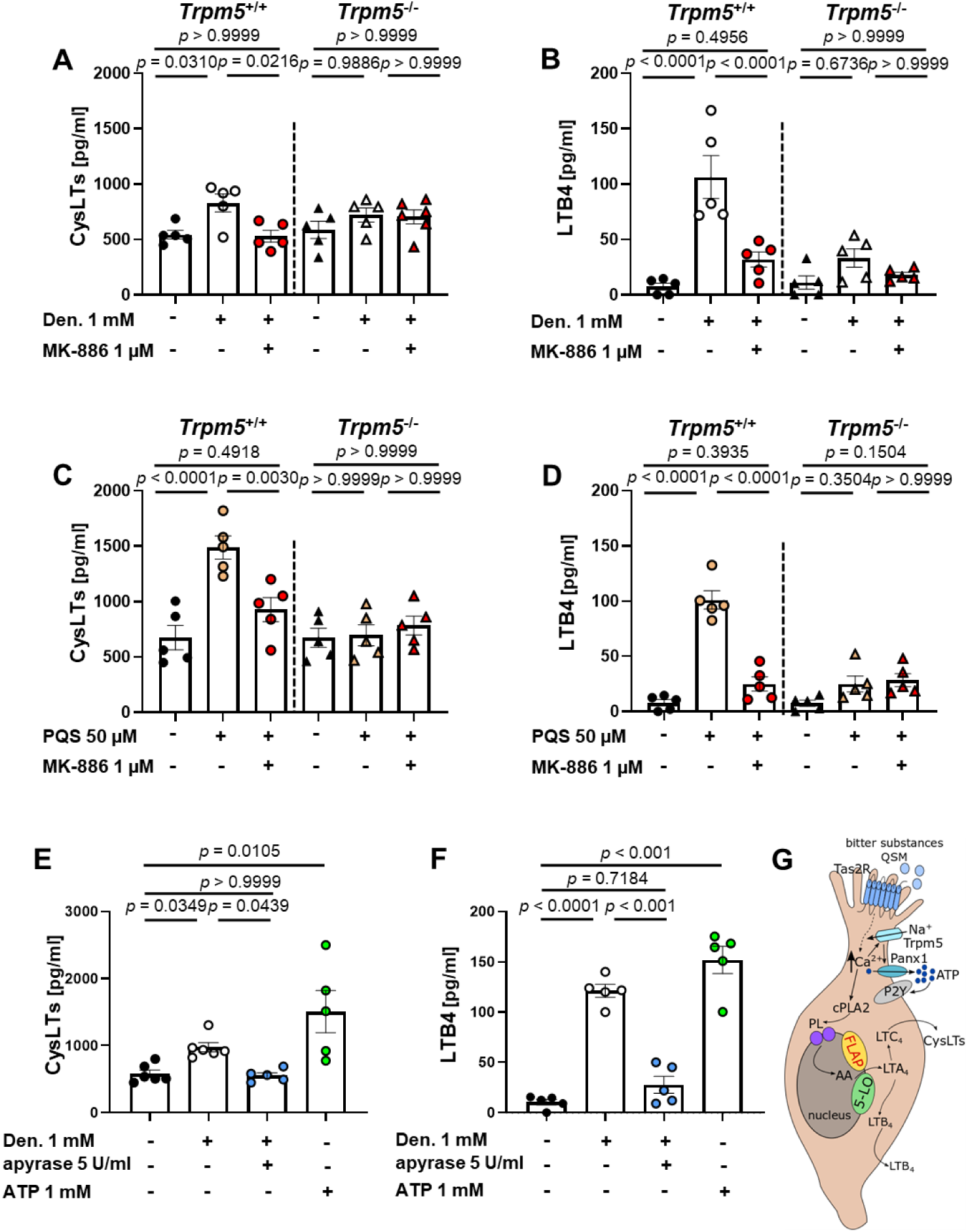
Tracheal tuft cell-mediated release of CysLTs and LTB4 involves activation of the canonical bitter signaling cascade. (A-F) ELISA measurements of CysLTs and LTB4 in tracheal supernatants. Tracheae of *Trpm5^+/+^* (n=5-6) and *Trpm5^-/-^* (n=5) mice were stimulated with denatonium (Den., 1 mM), *Pseudomonas aeruginosa* quinolone signal (PQS) (50 µM) or ATP (10 mM). Where indicated, tracheae were incubated with the FLAP-inhibitor MK-886 (1 µM) or with apyrase (5 U/ml). Data are presented as mean ± SEM of 5-6 independent experiments; each data point represents an individual animal. One-way ANOVA followed by Bonferroni’s multiple-comparison (A-D, and F) or Kruskal-Wallis with Dunn’s test for pairwise multiple comparisons (E). (G) Schematic drawing: Tas2R agonists, e.g., denatonium, trigger release of ATP from tuft cells, which in turn acts in an autocrine manner to induce the generation of CysLTs and LTB4.

### Tracheal tuft cells recruit neutrophils via LTB_4_ release

Trpm5-dependent tracheal tuft cell activation initiates immune responses mediated by sensory neurons leading to plasma extravasation and neutrophil recruitment^2^. These responses largely depend on the release of the neuropeptide calcitonin gene-related peptide (CGRP) from Trpa1^+^ sensory nerve terminals^2^. We next asked whether tuft cell-derived LTB_4_ acts on sensory neurons to induce CGRP release. We first confirmed the involvement of sensory neurons in tuft cell-mediated neurogenic inflammation using a resiniferatoxin (RTX) mouse model, in which Trpv1^+^ (and Trpa1^+^) neurons were selectively ablated^32^ (Figures S5A and B). Sensory neuron ablation abolished both plasma extravasation and neutrophil recruitment induced by tracheal tuft cell activation with denatonium (Figures S5C and D). The reduced number of neutrophils recruited to the trachea was accompanied by elevated neutrophil levels in the blood of RTX-treated mice (Figure S5E). Furthermore, sensory neuron ablation with RTX did not significantly affect the LTB_4_ release upon tuft cell activation (Figure S5F). Since LTs can activate sensory neurons and drive vascular changes associated with neurogenic inflammation^33–35^, we next investigated whether LTs released from activated tuft cells might contribute to these inflammatory responses. We blocked LT synthesis *in vivo* using MK-886 (10 mg/kg; *i.p.*) and evaluated the ability of tracheal tuft cells to induce plasma extravasation and neutrophil recruitment 30 min after intratracheal denatonium administration. Denatonium (1, 10, and 20 mM) induced a dose-dependent increase in plasma extravasation in both vehicle-treated and MK-886-treated *Trpm5^+/+^* mice (Figure 3A). Surprisingly, neutrophil recruitment to the lamina propria following denatonium stimulation was significantly reduced in *Trpm5^+/+^* mice treated with MK-886, compared to vehicle-treated controls (Figure 3B). Moreover, neutrophil numbers were elevated in blood from MK-886-treated *Trpm5^+/+^* mice compared to vehicle-treated controls (Figure S6A).

**Fig. 3:**
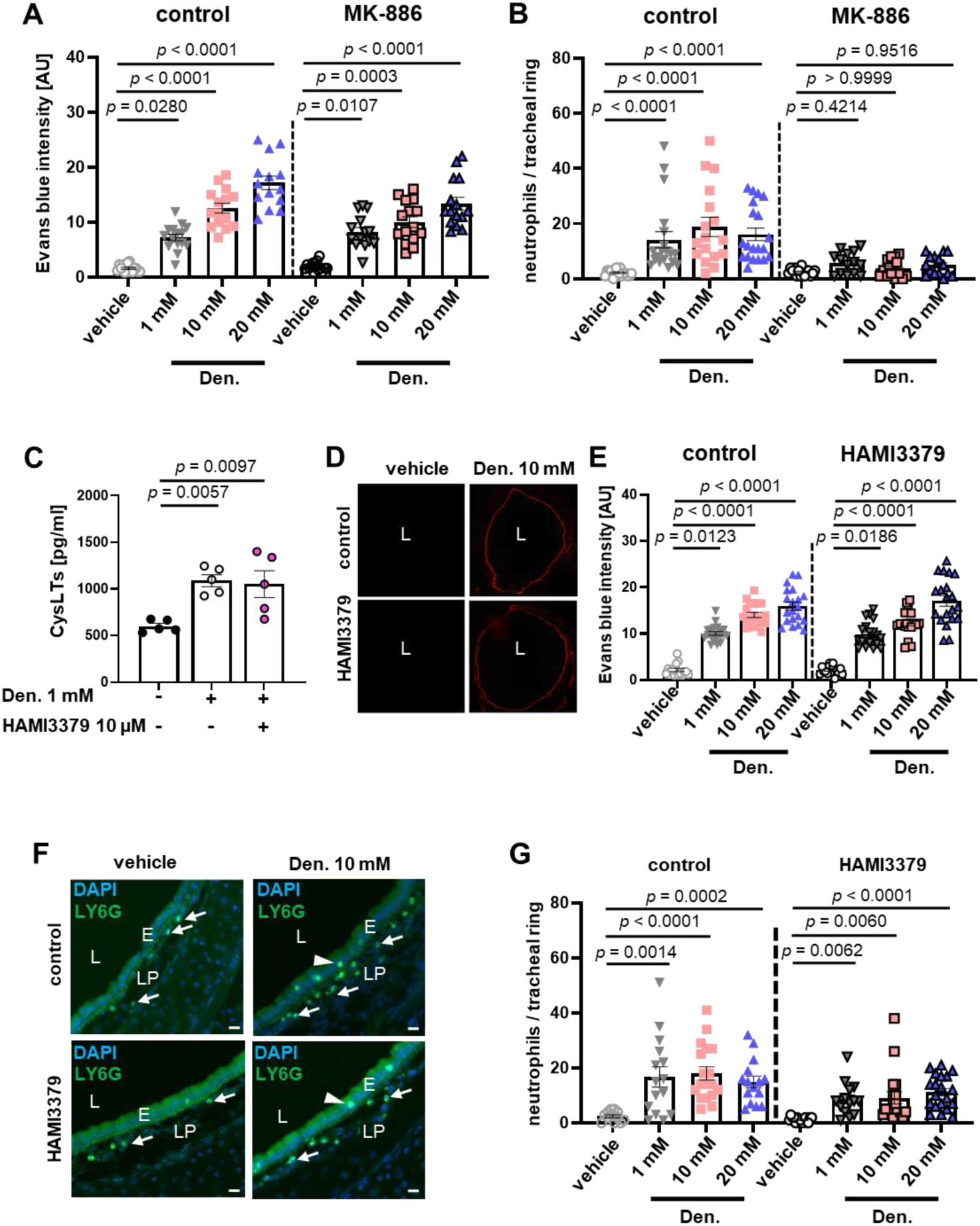
Activation of tracheal tuft cells triggers LTB4-induced neutrophil recruitment. (A) Evaluation of Evans blue in tracheal rings 30 minutes after intratracheal application of vehicle (PBS) or denatonium (Den.; 1 mM, 10 mM, 20 mM) in control (n=4) or MK-886-treated (10 mg/kg) (n=4) mice. (B) Quantification of recruited neutrophils in tracheal rings in response to vehicle or Den. in control (n=4) or MK-886 treated mice (10 mg/kg) (n=4). (C) ELISA measurements of CysLTs in tracheal supernatants in the presence of 10 µM HAMI3379 (n=5). (D) Representative images of tracheal rings showing EB fluorescence in response to vehicle and to 10 mM Den. in vehicle-(control) and HAMI3379-injected mice. (E) Quantification of EB in vehicle-(control) and HAMI36379-treated mice in response to Den. (n=4 mice, 15-20 tracheal rings). (F) Representative images from tracheal tissue sections stained for Ly6G showing neutrophils (arrows) in the lamina propria and the tracheal epithelium (arrowheads) in response to intratracheal administration of vehicle or 10 mM Den. in vehicle-(control) and HAMI36379-treated mice. Scale bar, 20 µM. L, lumen; E, epithelium; LP, lamina propria. (G) Quantification of the neutrophil numbers per tracheal ring in control and experimental animals in response to denatonium. Data are mean ± SEM of 12-20 tracheal rings of 4 mice (A, B, E, and G) and 5 mice (C). Kruskal-Wallis with Dunn’s test for pairwise multiple comparisons (A-B) or One-way ANOVA followed by Bonferroni’s multiple-comparison test (C, E and G); the p-value from the comparison between control and HAMI3379 groups treated with vehicle (PBS), 1, 10 and 20 mM denatonium are > 0.999, > 0.999, 0.0555, and > 0.999, respectively.

Previous studies have shown that sensory neurons of the dorsal root ganglia (DRG) and jugular-nodose ganglia (JNG), which innervate the airways, express CysLT receptor 2 (Cysltr2)^35–37^. We hypothesized that CysLTs released upon tuft cell activation might contribute to these inflammatory responses by activating the CysLTR2 on sensory neurons. To test this hypothesis, *Trpm5^+/+^* mice were injected with the selective CysLTR2 antagonist HAMI3379 (0.2 mg/kg; *i.p.*) on two consecutive days before intratracheal administration of denatonium. HAMI3379 did not alter the increased CysLTs levels triggered by tuft cell activation with 1 mM denatonium (Figure 3C). Moreover, inhibition of CysLTR2 did not affect the tuft cell-induced plasma extravasation (Figures 3D and E) and neutrophil recruitment, as no significant differences were observed between vehicle- or HAMI3379-treated mice following denatonium administration (Figures 3F and G). Similarly, no differences were detected in the percentage of blood neutrophils when the mice were injected with HAMI3379 (Figure S6B). We next considered the possibility that ILC2s might contribute to neurogenic inflammation, as CysLTs activate ILC2s in the lung and induce a release of interleukin 5 (IL-5)^38^. Since IL-5 might activate sensory neurons to release immune cell-modulating peptides^39,40^, we investigated the involvement of tuft cell-mediated ILC2-activation for the induction of a neurogenic inflammatory response using the *Il5^-/-^* mouse model. Unexpectedly, intratracheal denatonium application in the *Il5^-/-^* mice did neither affect plasma extravasation nor neutrophil recruitment in comparison to *Il5^+/+^* mice (Figures S7A-C). These results indicate that CysLTs released from tracheal tuft cells in response to Tas2R agonists are dispensable for initiating acute neurogenic inflammation.

Finally, we hypothesized that the release of CGRP is primarily required for the enhancement of the vascular permeability and the plasma extravasation^41^, while the release of LTB_4_, which is a potent chemotactic factor^42^, drives neutrophil recruitment to the site of inflammation. To assess the impact of tuft cell activation on the migration capacity of neutrophils, we performed transwell migration assays with primary mouse lung endothelial cells isolated by FACS-sorting (Figures S8A and B). Supernatants obtained from tracheal tuft cells stimulated with 1 mM denatonium increased neutrophil migration across the endothelial layer significantly (Figure 4A). This migration was mainly dependent on LTB_4_, as supernatants from tuft cells stimulated with denatonium reduced neutrophil migration to baseline levels in the presence of 1 µM MK-886 (Figure 4A). Consistent with this, LTB_4_ stimulated neutrophil migration to the same extent as supernatants from denatonium-stimulated tuft cells. In control experiments, the administration of MK-886 (10 mg/kg) alone did not affect the migration of neutrophils toward the potent chemoattractant N-formyl-l-methionyl-l-leucyl-l-phenylalanine (fMLF; 100 nM) (Figure S8C). Interestingly, there was no enhanced migration of neutrophils toward CGRP (1 ng/ml) (Figure 4A). To further assess vascular permeability, we also measured changes in transendothelial electrical resistance (TEER). TEER was reduced when endothelial cells were treated with supernatants obtained from tuft cells stimulated with 1 mM denatonium, indicating an increased permeability of the lung endothelial cell layer (Figure 4B). Similar results were obtained when 1 ng/ml CGRP was added to the cell culture medium. LTB_4_ did not affect TEER, and pharmacological inhibition of its synthesis in the trachea using MK-886 failed to significantly attenuate the denatonium-induced response (Figure 4B). Blocking CGRP receptors *in vivo* with the specific antagonist (CGRP_8-37_) resulted in elevated levels of circulating neutrophils after intratracheal application of denatonium (Figure 4C). Of note, inhibition of LT release by MK-886 did not affect the ability of tuft cells to induce CGRP release (Figure 4D). These findings indicate that the increase in vascular permeability and plasma extravasation following tuft cell activation primarily depends on CGRP. This increased permeability is a prerequisite for neutrophil transmigration at the site of inflammation, whereas LTB_4_ serves as a key chemotactic signal that especially attracts the neutrophils to the site of tuft cell activation (Figure 4E).

**Fig. 4:**
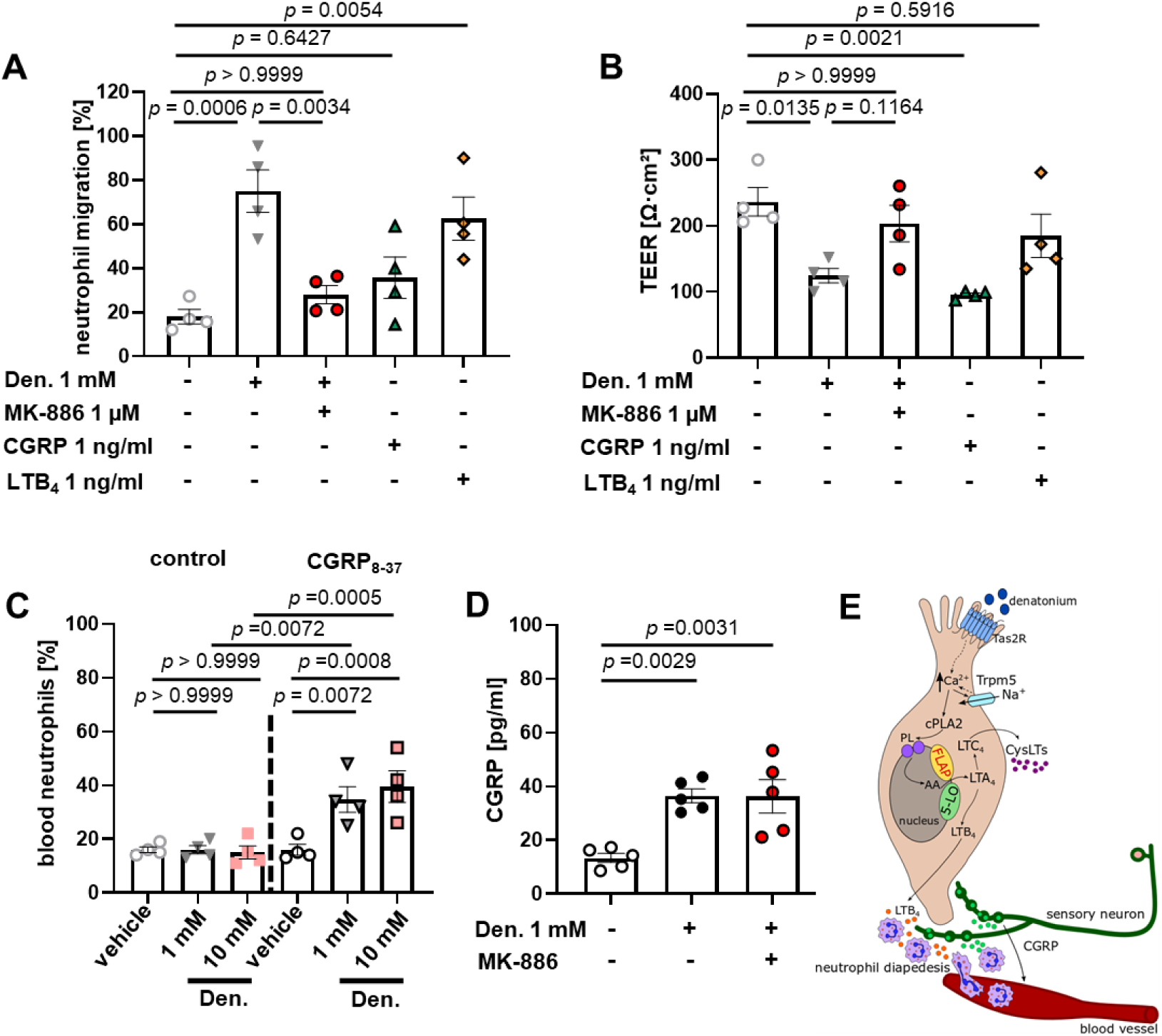
Neutrophil recruitment following tuft cell activation is mediated by LTB4. (A-B) Quantification of neutrophil migration across a lung endothelial cell layer in a transwell assay. Endothelial cells were treated with vehicle (PBS), denatonium (Den., 1 mM), denatonium plus MK-886 (1 µM), CGRP (1 ng/ml), or LTB4 (1 ng/ml). Data represent the percentage of migrating neutrophils (n=4). (B) Measurement of transendothelial electrical resistance (TEER) of the lung endothelial cell layer (n=4). (C) Analysis of blood neutrophils following blocking of CGRP with CGRP_8-37_ (n=4) (D) Measurement of CGRP by ELISA (n=5). (E) Activation of tuft cells induces LTB4 release, promoting neutrophil recruitment. Tuft cell-mediated CGRP release from nerve fibers leads to vasodilation and increased vascular permeability, and enables the neutrophil transmigration from dilated blood vessels. Data are shown as mean ± SEM of 4-5 independent experiments. One-way ANOVA followed by Bonferroni’s multiple-comparison test (A-D).

### Tracheal tuft cell-released LTs trigger early recruitment of neutrophils and macrophages in *P. aeruginosa* infection

Next, we tested the role of LTs produced by tracheal tuft cells in *P. aeruginosa* infection. Depletion of LT synthesis was achieved by the *i.p.* administration of MK-886 (10 mg/kg) 30 min before infection of the mice with a 4 h culture of the mucoid *P. aeruginosa* strain NH57388A grown in LB medium. Tracheae, BALF, blood, and spleens were collected for FACS analysis 4 h after infection. We found that total leukocyte, neutrophil, and macrophage counts significantly increased in tracheae (Figures 5A-C, gating strategies in Figure S9A) and monocyte counts in BALF (Figure 5D-F, gating strategies in Figure S9B) of infected *Trpm5^+/+^* mice but remained at basal levels in sham-treated and infected animals that received MK-886 prior to infection. In addition, we found that *P. aeruginosa* infection in mice led to LTB_4_ release in the BALF, which was dependent on *Trpm5* (Figure S10A). In contrast, total leukocyte and neutrophil counts in the blood of *P. aeruginosa*-infected mice did not differ from those of sham-treated controls but were strongly increased when LT synthesis was inhibited with MK-886 prior to infection (Figures S10B and C gating strategies in Figure S11A). Leukocyte, neutrophil, and macrophage counts remained unchanged in *Trpm5*^⁻/⁻^ mice, in contrast to the robust immune response observed in *Trpm5*⁺^/^⁺ mice (Figure 5), further corroborating the functional role of Trpm5 in tuft cell-induced immune responses. Moreover, we found a significant increase in neutrophil counts in the spleen of infected *Trpm5^-/-^* mice, pointing toward splenic sequestration of these cells (Figure S10D, gating strategies in Figure S11B). Together, these findings indicate that tuft cell activation and subsequent LT release are crucial for combating bacterial infection. The inhibition of LT signaling or impaired Trpm5-signaling in response to *P. aeruginosa* infection leads to an increased number of leukocytes in the blood, particularly of neutrophils, likely because they fail to transmigrate from the vasculature to the site where the pathogens are located and instead accumulate in the spleen.

**Fig. 5:**
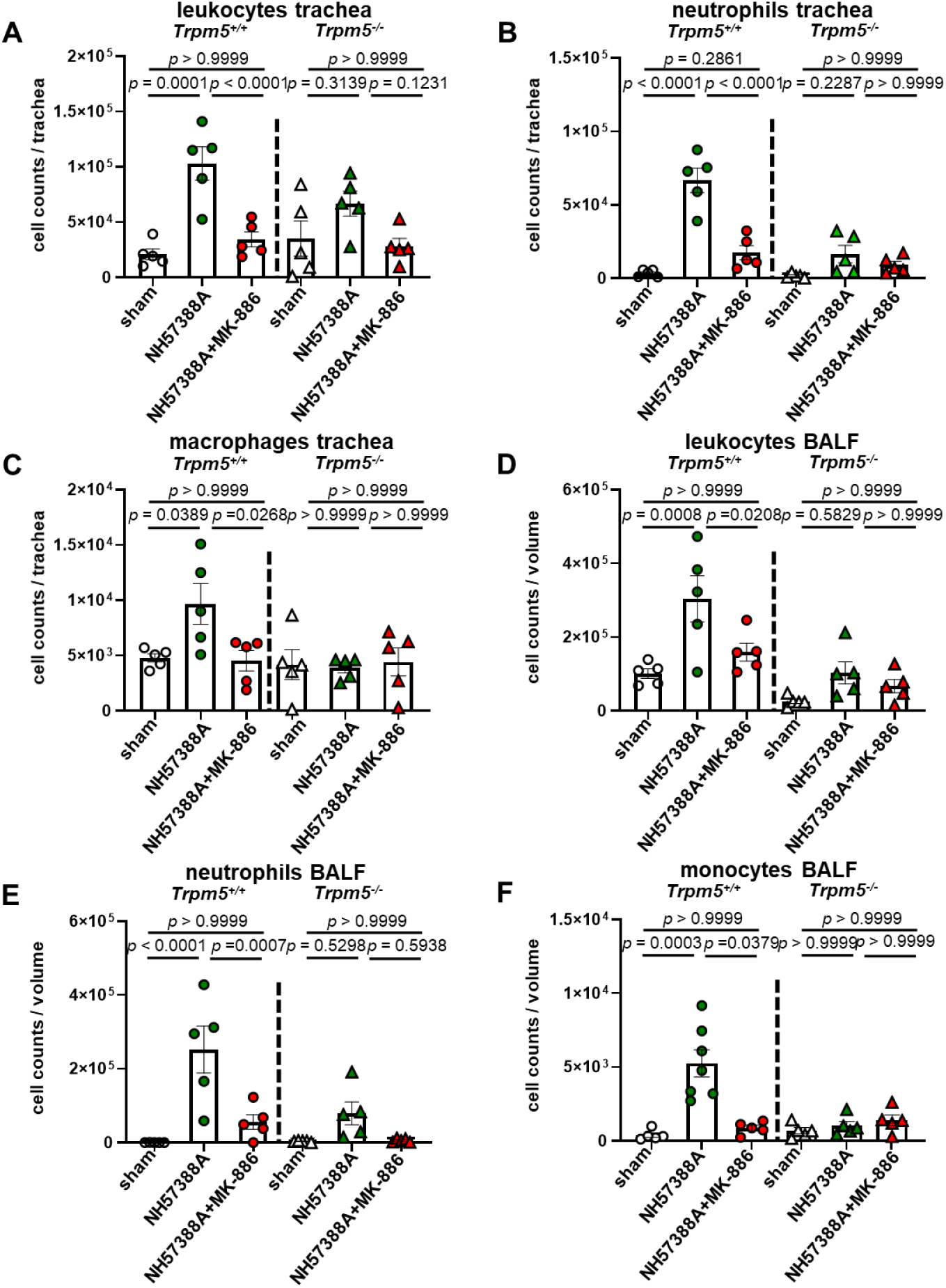
LTs contribute to the recruitment of immune cells in respiratory infection with *P. aeruginosa* NH57388A. (A-F) FACS analysis of leukocytes (CD45^+^), neutrophils, and macrophages/monocytes in the trachea and BALF of *Trpm5^+/+^* mice either sham-treated (n=5), infected with NH57388A (n=5) or infected with NH57388A following the administration of MK-886 (10 mg/kg) (n=5), or *Trpm5^-/-^* mice either sham-treated (n=5) or infected with NH57388A (n=5). Data are presented as mean ± SEM of 5 animals; each symbol represents an animal (A-F). One-way ANOVA followed by Bonferroni’s multiple-comparison test (A-E) or Kruskal-Wallis with Dunn’s test for pairwise multiple comparisons (F).

### Tracheal tuft cells elicit a rapid immune response to bacterial ATP

Previous studies have shown that clinical isolates from *P. aeruginosa*, among other bacterial pathogens, secrete ATP extracellularly^43,44^; however, the clinical relevance of the bacterial extracellular ATP (eATP) in respiratory infection is not well understood. Since tuft cells express a variety of purinergic receptors, including P2Y2, P2X4, and P2X6^14–16^, we hypothesized that they might act as ATP sensors during bacterial infections, and respond to elevated eATP levels through the release of LTs. To test our hypothesis, we first measured the eATP levels secreted by the *P. aeruginosa* isolate NH57388A at different time points of bacterial growth (Figures 6A and B). We observed that eATP levels in the bacterial cultures increased after 2 h of growth, reached a maximum at 4 h, and subsequently declined to undetectable levels at 12 h (Figure 6B). In contrast, the QSM levels, including PQS (Figure 6C), 2-aminoacetophenone (2AA), 2-heptylhydroxyquinoline N-oxide (HQNQ), 4-hydroxy-2-heptylquinoline (HHQ), 2,4-dihydroxyquinoline (DHQ), and N-Butanoyl-L-homoserine lactone (C4-HSL) increased at later stages of growth for all except DHQ levels, which exhibited a slow upward trend after 4 h of growth (Figures S12A-E). These QSMs are released in later stages of bacterial growth and activate tracheal tuft cells^2^. Additionally, we confirmed that other pathogenic respiratory bacteria, including the *S. pneumoniae* strain PN36, *K. pneumonia* strain Kp52145, and *S. aureus* strain Newman, release ATP during the early phase of their growth (Figures S13A-F). Based on these observations that 4 h of growth represented the time point at which *P. aeruginosa* supernatants contained the highest concentration of ATP and no detectable QSMs, we collected these supernatants to investigate the impact of bacterial eATP on tuft cell-induced immune responses. When tracheae were exposed to these supernatants, we found a Trpm5-dependent LTB_4_ and CysLT release in tracheal supernatants, since this was abolished in *Trpm5*^+/+^ by MK-886 and was not detected in *Trpm5*^-/-^ mice (Figures 6D and E). We next asked whether activation of tracheal tuft cells with bacterial supernatants could induce acute immune responses. To test this, we treated *Trpm5*^+/+^ and *Trpm5*^-/-^ mice intratracheally with bacterial supernatants collected after 4 h of growth and analyzed the induction of neutrophil recruitment in the trachea. We found an increased recruitment of neutrophils to the trachea in *Trpm5*^+/+^ but not in *Trpm5*^-/-^ mice (Figure 6F). Moreover, analysis of neutrophil percentages in the blood revealed an increased number of circulating neutrophils in *Trpm5*^-/-^ mice, whereas no such increase was observed in *Trpm5*^+/+^ mice (Figure 6G). Bacterial supernatants pretreated with apyrase (5 U/ml) to degrade eATP (Figure S14A) failed to stimulate LTB_4_ release from tracheal tuft cells (Figure S14B) as well as the subsequent neutrophil recruitment to the tracheal lamina propria *in vivo* (Figures S14C). Together, these findings indicate that LTB_4_ is released from tracheal tuft cells *in vivo* upon stimulation with bacterial eATP and is crucial for the recruitment of neutrophils during bacterial infections. To assess how tuft cell-released LTB_4_ shapes immune responses to *P. aeruginosa* induced *in vivo*, we analyzed the composition of the immune cell populations in the BALF of *Trpm5^-/-^* and *Trpm5^+/+^* mice after tracheal inhalation of 4 h supernatants. We observed elevated leukocyte, neutrophil, and mononuclear cell (MNCs) numbers - e.g., monocytes, macrophages, and lymphocytes - in *Trpm5^+/+^* mice, whereas this increase was absent in *Trpm5^-/-^* mice (Figures 7A-F). This enhanced immune cell recruitment depended on bacterial eATP and was mediated by LTs released from tuft cells, as blocking of LT synthesis with MK-886 in *Trpm5^+/+^* mice abolished this response (Figures 7A-F). Additionally, we observed an increased expression of the activation markers CD80 and CD86 in BALF monocytes only in *Trpm5^+/+^* mice and not in their *Trpm5^-/-^* littermates. This enhanced expression was LT-dependent, since administration of MK-886 (10 mg/kg) restored expression to basal levels (Figures S15A and B).

**Fig. 6:**
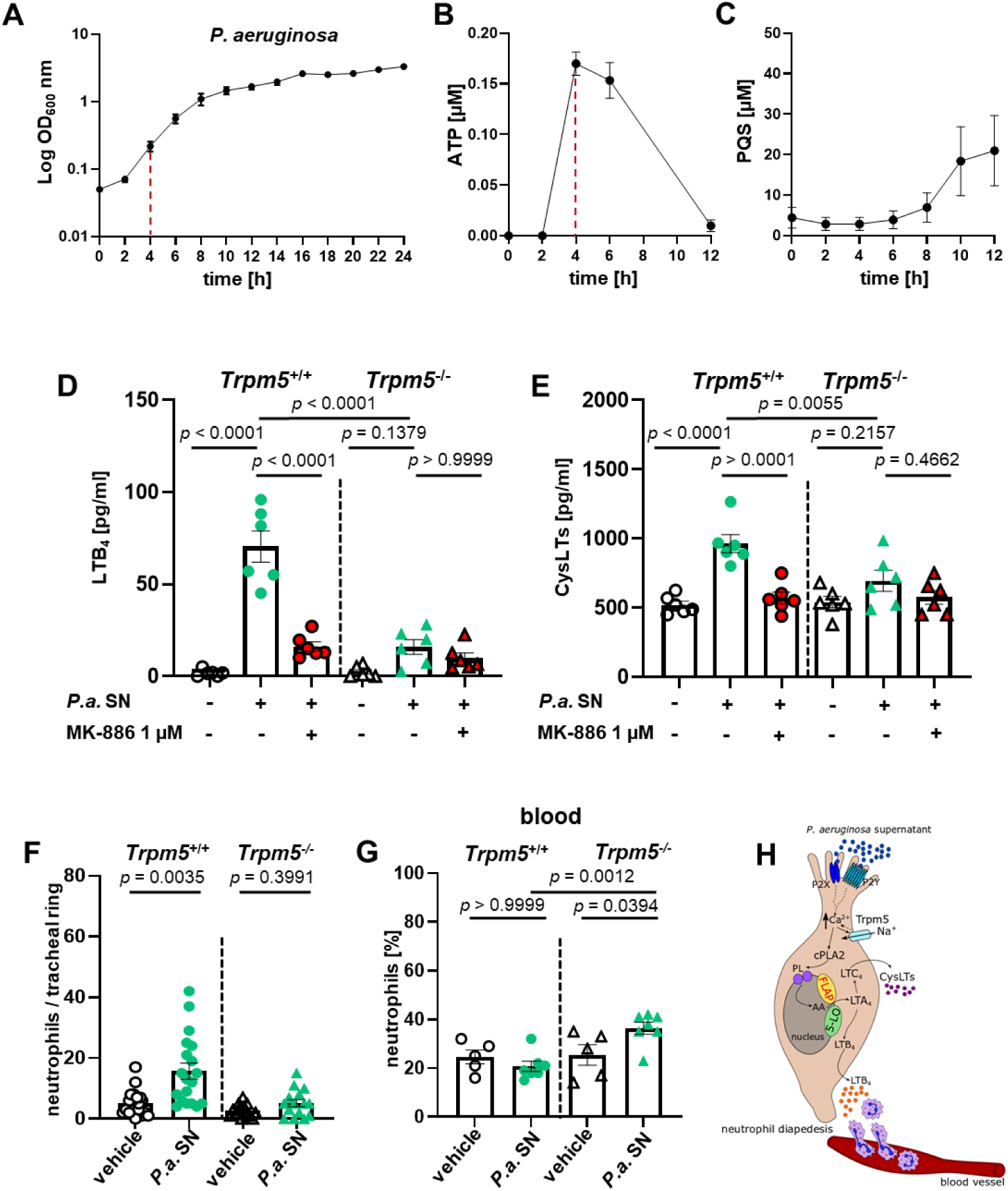
Bacterial ATP triggers neutrophil recruitment in a Trpm5-dependent manner. (A) Growth curve of the *P. aeuroginosa* isolate NH57388A (n=5). (B) Measurements of ATP released at different time points during cultivation of the *P. aeuroginosa* strain NH57388A using BacTiter-Glo™ Microbial Cell Viability Assay Reagent (n=3). (C) Mass-spectrometry measurements of PQS at different time points of cultivation of *P. aeuroginosa* strain NH57388A (n=5). (D) ELISA measurements of LTB_4_ in tracheal supernatants of *Trpm5^+/+^* (n=6) or *Trpm5^-/-^* tracheae (n=6) treated with either vehicle, NH57388A supernatant (*P.a.* SN) collected after 4h of growth, or *P.a.* SN and MK-886 (1 µM). (E) Quantification of CysLT release from *Trpm5^+/+^* (n=6) or *Trpm5^-/-^* tracheae (n=6) exposed the same experimental conditions as in D. (F) Quantification neutrophil infiltration in the tracheal lamina propria in *Trpm5^+/+^* (n=4) and *Trpm5^-/-^* (n=4) of tracheal sections 30 min after intratracheal administration of vehicle (controls) or NH57388A supernatants collected from 4h *P. aeruginosa* cultures (*P.a.* SN). (G) Percentages of neutrophils in the blood of mice (n=5-7) 30 min after intratracheal administration of NH57388A supernatants used in F. (H) Tracheal tuft cells induce a recruitment of neutrophils in a Trpm5-dependent manner in response to bacteria-secreted ATP. Data are shown as mean ± SEM of 3-5 biological replicates with each sample measured in duplicate (A to C), 6 mice (D-E), 15-20 tracheal rings of 4 mice (F), and 5-7 mice (G). One-way ANOVA followed by Bonferroni’s multiple-comparison test (D-H).

**Fig. 7:**
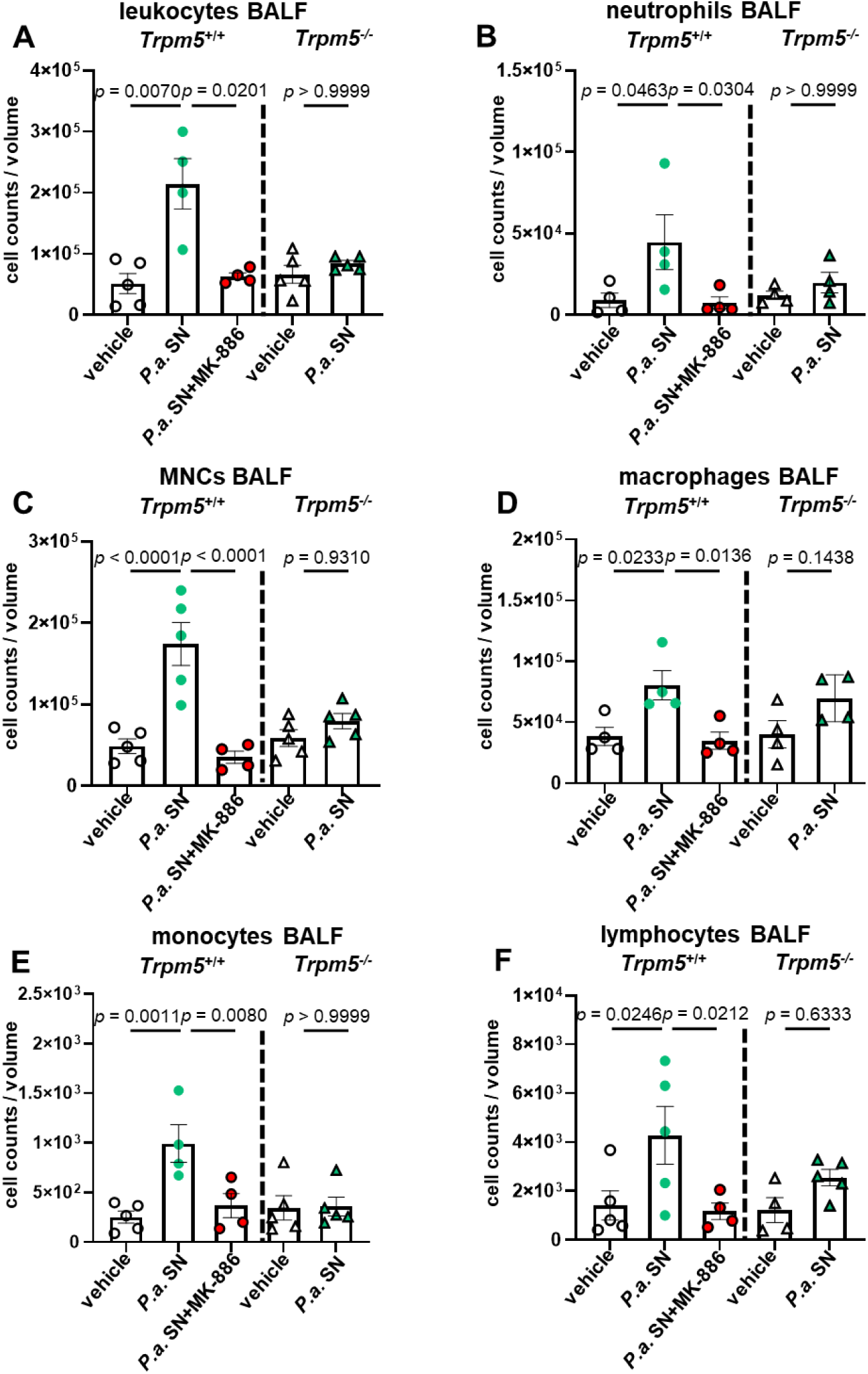
Bacterial supernatants from *P. aeruginosa* NH57388A alter the immune cell populations in BALF in a Trpm5- and LT-dependent manner. **(A-F)** Cell counts of leukocytes, neutrophils, and mononuclear cells (MNCs) in BALF from *Trpm5^+/+^* (n=4-5) and *Trpm5^-/-^* mice (n=4-5), 30 min after intratracheal application of either vehicle (medium) or NH57388A supernatants (*P.a.* SN, 4 h). Data represent mean ± SEM of 4-5 mice. One-way ANOVA followed by Bonferroni’s multiple-comparison test (A and C-F) or Kruskal-Wallis with Dunn’s test for pairwise multiple comparisons (B).

### Tuft cells modulate the bactericidal activity of airway macrophages

Next, we investigated the impact of tuft cell-derived LTs on the bacterial killing capacity of airway macrophages. Tracheal and lung interstitial macrophages were isolated from *Trpm5^+/+^* mice using FACS (Gating strategies in Figure 10A, and Figure S16) and cultured for 30 min with supernatants obtained from *Trpm5*^+/+^ or *Trpm5*^-/-^ mouse tracheae stimulated with 1 mM denatonium. Both tracheal and lung interstitial macrophages showed enhanced bacterial phagocytic capacities at 1 h after infection with *P. aeruginosa* strain NH57388A when they were exposed to supernatants obtained from denatonium-treated *Trpm5*^+/+^ tracheae prior to infection (Figures 8A-C). Exposure of the lung interstitial macrophages to *Trpm5*^-/-^ tracheal supernatants did not affect their bacterial killing capacity (Figure 8B-C). Specific tuft cell stimulation with CNO in the *Trpm5-*DREADD mouse model resulted in an enhanced bacterial killing capacity of lung interstitial macrophages (Figure 8D). To explore the role of tuft cell-derived LTs in macrophage-mediated bacterial killing of *P. aeruginosa*, we inhibited the synthesis of LTs by incubating the trachea with CNO in the presence of 1 µM MK-886. Macrophage killing capacity was significantly reduced compared to that of macrophages treated with supernatants from stimulated tuft cells without inhibition of LT synthesis and was comparable to baseline activity (Figure 8D). Next, we investigated the effect of the T2R agonist denatonium on the killing capacity of murine lung interstitial macrophages to exclude the possibility that their activation was due to a direct stimulation of T2Rs expressed in human macrophages^45^. Indeed, incubation of the murine interstitial macrophages with 1 mM denatonium did not increase their capacity to kill the *P. aeuroginosa* isolate NH57388A (Figure S17A). We considered the CysLT receptor as a potential mediator of the enhanced bacterial killing activity of macrophages following treatment with tracheal supernatants obtained after tuft cell activation. We found that lung interstitial macrophages expressed both *Cysltr1* and *Cysltr2*, as well as *Ltb4r1*, but did not express *Ltb4r2* (Figure 8E). Inhibition of the detected receptors by montelukast for Cysltr1 (1µM), but not HAMI3379 (10 nM) for Cysltr2 or CP105696 (1 µM) for Ltb4r1, in macrophages treated with tracheal supernatants collected after tuft cell activation with denatonium and infected with the *P. aeuroginosa* strain NH57388A reduced their bacterial killing capacity (Figure 8F). This finding indicates that tuft cell-released CysLTs activate lung interstitial macrophages through CysLTRs to enhance their bactericidal activity. To mechanistically assess how tuft cell-induced macrophage activation leads to *P. aeruginosa* clearance, we studied the release of hydrogen peroxide (H_2_O_2_), which has been previously shown to be detrimental to bacterial cells^46,47^. For this, we treated macrophages with tracheal supernatants obtained from *Trpm5^+/+^* and *Trpm5^-/-^* mice. Activation of macrophages with tracheal supernatants obtained from *Trpm5^+/+^* tracheae stimulated with denatonium induced a robust production of H_2_O_2_ (Figure 8G), whereas supernatant from *Trpm5^-/-^* tracheae had no effect, indicating that the observed H_2_O_2_ release was dependent on tuft cell activation and signaling. Moreover, we found that H_2_O_2_ production by macrophages was induced by LTs released upon activation of tracheal tuft cells (Figure 8H). The enhanced release of H_2_O_2_ was associated with an increased expression of *Nox2* (encoding NADPH oxidase 2) in macrophages in a LT-dependent manner (Figure S17B).

**Fig. 8:**
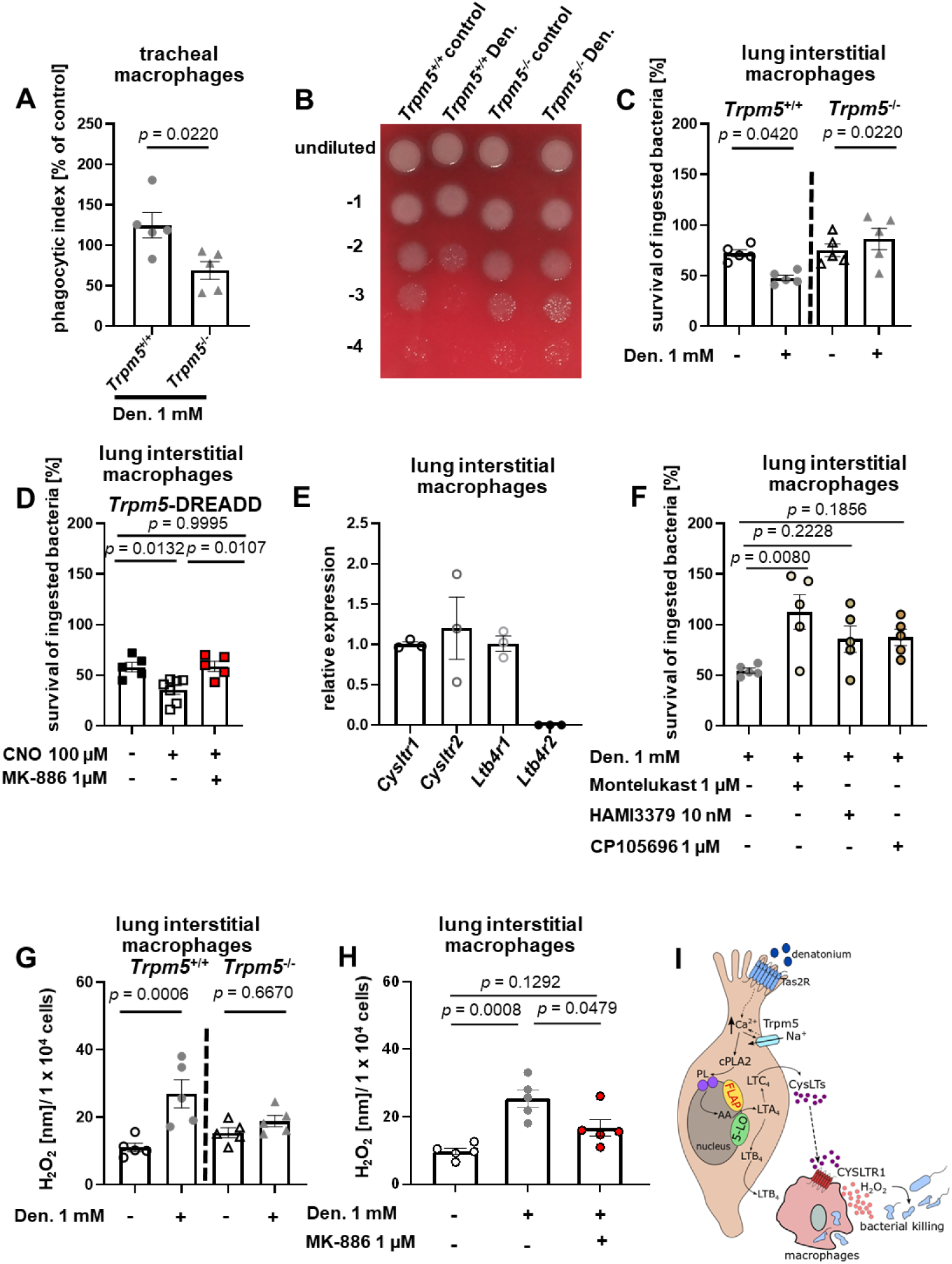
Trpm5 signaling increased the bacterial killing capacity of tracheal and lung interstitial macrophages. (A) Bacterial killing assay with tracheal macrophages primed with tracheal supernatants from explanted *Trpm5^+/+^* (n=5) and *Trpm5^-/-^* (n=5) tracheae following stimulation with denatonium (Den., 1 mM). (B) Representative image of a blood agar plate showing *P. aeruginosa* NH57388A survival after phagocytosis by lung interstitial macrophages activated with tracheal supernatants from *Trpm5*^⁺/⁺^ or *Trpm5*^⁻/⁻^ tracheae stimulated with vehicle or 1 mM denatonium (Den). (C-D) Bacterial killing assay with lung interstitial macrophages stimulated with supernatants from explanted tracheae from *Trpm5^+/+^* (n=5) or *Trpm5^-/-^* (n=5) mice treated with vehicle or denatonium (C), and from *Trpm5*-DREADD tracheae (n=5-7) incubated with CNO (100 µM) or CNO and MK-886 (1 µM) (D). (E) qRT-PCR analysis of *Cysltr1*, *Cysltr2*, *Ltb4r1,* and *Ltb4r2* in lung interstitial macrophages depicted as relative gene expression (n=3). (F) Bacterial killing assay with lung interstitial macrophages stimulated with supernatants from explanted *Trpm5^+/+^* mouse (n=5) tracheae treated with Den., either alone or in combination with either Montelukast (1 µM), HAMI3379 (10 nM), or CP105696 (1 µM). (G-H) Quantification of H_2_O_2_ released upon activation of lung interstitial macrophages with tracheal supernatants of *Trpm5^+/+^* (n=5) and *Trpm5^-/-^* mice (n=5) treated with vehicle or Den., or of *Trpm5^+/+^* tracheae treated with vehicle, Den., or Den. with MK-886 (1 µM). (I) Tuft cell-released CysLTs enhance the bacterial killing capacity of airway macrophages by inducing a release of H_2_O_2_. Data represent mean ± SEM of 3-7 independent experiments. One-way ANOVA followed by Bonferroni’s multiple-comparison test.

### The tuft cell-sensing capacity of bacterial ATP is required to combat bacterial infections

The ability of tracheal tuft cells to detect bacterial eATP resulted in enhanced recruitment of immune cells to the trachea and alveolar spaces. Thus, we postulated that tuft cells can differentiate between pathogenic and commensal bacteria in the airways and that this function is fundamental for pathogen clearance during infection and thus for maintaining microbiota homeostasis in the lower respiratory tract. Consistent with this hypothesis, *R. pneumotropicus* caused chronic respiratory infection in *Trpm5^-/-^*, but not in *Trpm5^+/+^* mice. 60% of naturally infected *Trpm5^+/+^* mice cleared the infection, whereas only 15.6% of infected *Trpm5^-/-^* mice showed recovery (Figure 9A). In addition, we isolated several commensal bacteria, including *Micrococcus luteus*, *Rothia nasimurium*, *Streptococcus mitis*, and *Staphylococcus xylosus,* from both *Trpm5^+/+^* and *Trpm5^-/-^* mice. Interestingly, none of these commensals released ATP during growth, in contrast to the pathogenic bacterium *R. pneumotropicus* (Figure 9B). Analysis of the immune cell populations revealed increased counts of neutrophils in the trachea of *Trpm5^+/+^* infected animals (Figure 9C). Furthermore, more leukocytes, neutrophils, and macrophages were recruited in the BALF of these mice (Figures 9D-F), whereas *Trpm5^-/-^* animals showed increased counts of neutrophils and monocytes in the blood (Figures 9G and H). These findings suggest that bacterial ATP functions as a virulence factor that influences bacterial colonization of the trachea. However, the presence of tracheal tuft cells with a functional Trpm5 signal transduction cascade is required to induce innate immune responses against these pathogens.

**Fig. 9:**
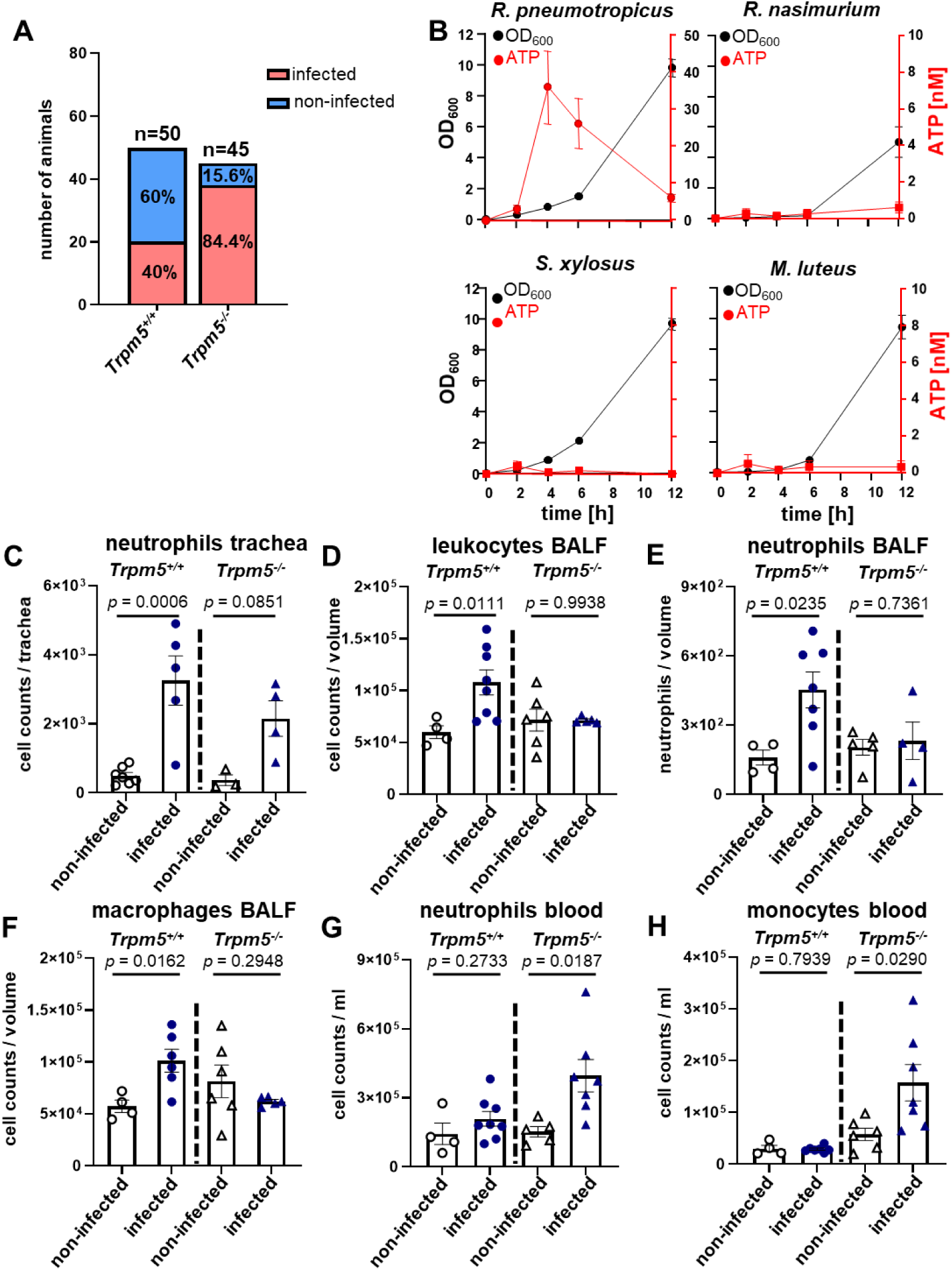
Mice with functional Trpm5 showed enhanced resistance against natural infections. (A) Percentages of *Trpm5^+/+^* (n=50) and *Trpm5^-/-^* (n=45) mice infected with *Rodentibacter pneumotropicus*. (B) Growth curves and measurement of extracellular ATP released by the pathogen *Rodentibacter pneumotropicus* and by the airway commensals *Rothia nasimurium*, *Staphylococcus xylosus, and Micrococcus luteus (*n=3). (C) FACS analysis of neutrophil numbers in the tracheae of *Trpm5^+/+^* (n=5-7) and *Trpm5^-/-^* (n=3-4) mice naturally infected with *R. pneumotropicus* or in non-infected controls. (D-F) FACS analyses of cell counts of leukocytes (CD45^+^), neutrophils, and macrophages in the BALF from naturally infected *Trpm5^+/+^* (n=4-8) and *Trpm5^-/-^* (n=4-6) mice with *R. pneumotropicus*, as well as from non-infected control mice. (G-H) FACS analyses of neutrophil and monocyte cell counts in the blood of *Trpm5^+/+^* (n=4-8) or *Trpm5^-/-^* (n=5-7) mice either naturally infected with *R. pneumotropicus* or non-infected. Data represent: (A) the number of animals per group (control or naturally infected mice), (B) mean ± SEM of 3 independent experiments performed in duplicates, and (C-H) mean ± SEM from 3-8 mice per group. One-way ANOVA followed by Bonferroni’s multiple-comparison test (C-H).

## DISCUSSION

Our work delineates the role of tracheal tuft cell-released LTs in the acute phases of bacterial pneumonia, especially for recruiting neutrophils and macrophages in the airways and alveolar space. We identify tuft cells as the primary source of proinflammatory LTs which are rapidly released in the presence of pathogenic bacteria in the airways. Mechanistically, we show that tracheal tuft cell activation with Tas2R agonists such as denatonium or bacterial PQS lead to LT release in a Trpm5-dependent manner. So far, CysLT production has been reported for intestinal tuft cells and nasal SCCs, although they are activated by different ligands (e.g. intestinal parasites versus aeroallergens) in these tissues^17,18,48^. We expand these observations and demonstrate that Trpm5 is involved in the generation of LTs from tracheal tuft cells in the context of bacterial pneumonia. We show that tuft cells discriminate between the presence of commensal and pathogenic bacteria in the airways and utilize LT-signaling to mediate pathogen clearance.

We have recently demonstrated that denatonium activation of tracheal tuft cells triggers ATP release^3^. Here, we address the hypothesis that tuft cell-derived ATP acts in an autocrine manner to induce LT synthesis and release. Such a scenario appears plausible since the nasal administration of ATP leads to CysLT-production by nasal SCCs^17^. Additionally, macrophages and astrocytes produce LTB_4_ and CysLTs, respectively, in response to eATP^49,50^. Denatonium tuft cell stimulation in the presence of the ATP-hydrolyzing enzyme apyrase inhibited LT release, indicating that ATP is required for LT production by tuft cells. We propose that the LT release by tuft cells involves a self-amplifying loop (reinforcing auto feedback loop), where Trpm5-dependent ATP release activates purinergic receptors, most likely P2Y2^17^, thereby increasing intracellular Ca^+2^ levels and activating LT-synthesizing pathways. We recently showed that activation of tuft cells with Ts2R agonists, such as denatonium or *P. aeruginosa* QSMs, induces a protective neurogenic inflammation characterized by plasma extravasation and neutrophil recruitment. These responses were mediated by the neuropeptide CGRP released from sensory nerve endings following tuft cell activation^2^. Cholinergic receptor inhibition abolished the induction of neurogenic inflammation. Here, we expand these findings and demonstrate that tuft cell-induced neurogenic inflammation occurs in two distinct phases and requires LT signaling. Building on our earlier findings that CGRP mediates neurogenic inflammation in response to bacterial and Tas2R stimulation, we now show that tuft cell-derived LTB_4_ contributes to this by promoting neutrophil recruitment, whereas CGRP primarily drives vasodilatation. Previous studies have shown that LTB_4_ activates neutrophils^42^ and promotes their recruitment to the lungs^51,52^. CGRP has previously been reported to increase the blood flow and the vascular permeability in the trachea and the knee joint^53,54^. Here, we propose a direct chemotactic effect for neutrophils to be mediated by the tuft cell-released LTB_4_. We excluded the involvement of sensory neurons in neutrophil recruitment. A subset of Trpv1^+^ DRG neurons expresses BLT1, the receptor for LTB_4_, which mediates calcium flux in response to ligand activation, and thereby induces neuropeptide release from these neurons^55,56^. Since LT synthesis inhibition with MK-884 did not affect CGRP release in the trachea, we assume that tuft cell-released LTB_4_ did not act on sensory nerve terminals located in the tracheal mucosa. Despite its expression in Trpv1^+^ sensory DRG neurons, CysLTR2 was also ruled out^35,37^ as a contributor to the tuft cell-driven neurogenic inflammatory response, as blocking CysLTR2 with HAMI3379 had no effect on neutrophil recruitment. Of note, DRG sensory neurons do not express CysLTR1^35^, indicating that CysLTs are not involved in tuft cell-driven neuronal activation. Nevertheless, tuft cell CysLTs released in response to *P. aeruginosa* might activate macrophages. Additionally, SP released from tuft cell-mediated and ACh-dependent stimulation of Trpa1+ sensory nerve fibers might also contribute to macrophage activation^2^. Activated macrophages could also secrete LTB_4_^57^ and contribute to the activation and chemotaxis of polymorphonuclear leukocytes (PMNs), such as neutrophils, to the trachea and BALF. Nevertheless, depletion of Trpv1^+^ sensory neurons with RTX abolished the acute extravasation of neutrophils following an *in vivo* tuft cell stimulation with denatonium or *P. aeruginosa* supernatants. CGRP receptor inhibition resulted in elevated neutrophil counts in circulation. RTX treatment abolished Evans blue extravasation, indicating that CGRP is responsible for the increased vascular permeability. The elevated number of neutrophils in the blood of RTX-treated mice following tuft cell stimulation further highlights the role of LTB_4_ in neutrophil recruitment^58^. These results point to a multifaceted response to tracheal tuft cell activation, involving both neurogenic signaling alongside the LTB_4_-mediated neutrophil recruitment.

We found that tracheal tuft cells function as sensors for eATP of bacterial origin. Although ATP released from damaged cells at later stages of infections is a well-established damage-associated molecular pattern (DAMP)^59^, our finding that ATP released by pathogenic bacteria upon growth is sensed by tracheal tuft cells highlights a fundamentally different mode of immune detection. eATP is secreted by bacterial pathogens early during *E. coli* and *Salmonella* infection^43,44^. We found that eATP production in the first 4 h of growth is a common feature of multiple respiratory pathogenic bacteria, such as *P. aeruginosa, S. pneumoniae, K. pneumoniae, S. aureus,* and *R. pneumotropicus*. Tuft cells sense bacterial eATP and rapidly induce immune responses in the presence of pathogens. Neutrophil recruitment and macrophage activation were LT- and Trpm5-dependent. Interestingly, in a mouse model of abdominal *E. coli*-induced sepsis, bacterial ATP reduced neutrophil counts, activated the endolysosomal system, and upregulated neutrophil degranulation, which together increased the severity of abdominal infection and early sepsis^60^. The decreased neutrophil counts can be explained by the rapid consumption of neutrophils in the face of sepsis, indicated by the better survival of mice in the absence of bacterial ATP. In this study, bacterial ATP release resulted from impaired membrane integrity and bacterial death. Other studies did not identify bacterial death as a relevant source of extracellular ATP during exponential growth^43^. It is plausible that ATP from living bacteria acts locally, while ATP released upon bacterial death via membrane vesicles acts systemically. Infection of mice with a 4 h culture of *P. aeruginosa*, which produces significant amounts of bacterial eATP, induced the recruitment of neutrophils and macrophages to the trachea and alveolar spaces. Thus, tuft cells act as sensors of bacterial eATP, enabling the detection of living bacteria or their metabolites early during infection, before significant tissue damage occurs. This unique detection strategy contrasts with the purely reactive DAMP-mediated response, which is typically triggered by initial tissue damage^61^. Trpm5-mediated migration of neutrophils is of crucial importance in infection^2^. Our data show that mice with defective Trpm5-signaling exhibit neutrophil sequestration in the spleen upon infection. Notably, splenic sequestration of neutrophils has been associated with systemic infections^62,63^. The ability of tuft cells to detect pathogenic bacterial eATP serves as a gatekeeper of microbial homeostasis in the airways and lungs. Colonization of airways with commensal bacteria which do not release eATP such as *M. luteus*, *R*. *nasimurium*, *S*. *mitis*, and *S*. *xylosus* did not trigger immune responses. The ATP-sensing capacity of tracheal tuft cells was critical for preventing colonization of the airways with the opportunistic bacterium *P. pneumotropica*. The pathogen resistance was accompanied by Trpm5-mediated recruitment of neutrophils and macrophages in the trachea of naturally infected mice. In *Trmp5^+/+^* mice, macrophages exhibited stronger antibacterial capacities and contributed to the clearance of the bacteria, ultimately leading to faster recovery from the *P. pneumotropica* infection. *Trpm5^-/-^* mice showed retention of macrophages and neutrophils in the blood and developed a chronic infection. Since the fundamental challenge for the host immunity at mucosal surfaces is to sustain tolerance between commensal bacteria while evoking robust immune responses against pathogens, our findings reveal sophisticated ATP-dependent mechanisms by which tracheal tuft cells act as tunable sensors differentiating between pathogenic and commensal bacteria.

Although our findings demonstrate a role of LTs in shaping tuft cell-mediated immune responses in the early stages of bacterial infection, several questions remain to be addressed in future studies. Since CysLTR3 is expressed by the airway epithelium^16^, it remains to be determined whether the tuft cell-derived LTs modulate the epithelial barrier function or the antimicrobial peptide release from neighboring epithelial cells. We recently showed that tuft cell activation facilitates the transition from innate to adaptive immunity by promoting Th17 cell recruitment^3^. Since LTs activate Th17 cells and induce their migration in the brain tissue^64,65^, further studies are needed to investigate whether the tuft cell-released LTs are involved in the Th17 cell migration. It is possible that Th17 cell activation following tuft cell stimulation occurs in a biphasic manner, in which the ATP released from tuft cells is followed by LT production, which promotes the migration of Th17 cells. Our findings uncover a previously unrecognized antibacterial function of tracheal tuft cells, identifying them as a key epithelial source of LTs that shape early immune responses in bacterial lung infection.

## METHODS

### Experimental animal models

All animal experiments and care procedures were approved by the animal welfare committee of Saarland (approval numbers 25/2021, 01/2023, and 21/2023) and were conducted according to the German guidelines for the care and use of laboratory animals. For the experiments, we used mice of both genders older than 8 weeks with the following genetic backgrounds: wild-type control animals, ChATBAC-eGFP (B6.Cg-Tg(RP23-268L19-EGFP)2Mik/J)^66^, *Trpm5*-DREADD^67^, *Trpm5*-DREADD-tGFP^67^, *Trpm5*^-/-^ (tuft cells deficient for Trpm5-signaling)^68^, *Il5^-/-^* (JAX Strain #:030926) and*Trpm5*-DTA (*Trpm5*-IRES-Cre-R26:*lacZbpA*^flox^DTA) which were generated by breeding by *Trpm*-5IRES-Cre mice^69^ with R26:*lacZbpA*^flox^DTA mice^70^. All lines were kept in a mixed (129/SvJ and C57BL/6J) background. The *ChAT^BAC^*-eGFP mice were bred and housed in the animal facility of the Institute of Experimental Surgery of the Saarland University in IVC cages. All other mice were bred and housed in the animal facility of the Institute of Experimental and Clinical Pharmacology of Saarland University under SPF conditions. All mice were kept under standardized 12 h day-night cycles with a humidity of 55 ± 10% and an ambient temperature of 22 ± 2 °C. Water and standardized food were provided *ad libitum*. No gender specific effects were investigated in this study. According to the respective protocol used for each experiment, mice were euthanized with an overdose of ketamine (Zoetis, Berlin, Germany)/xylazine (Bayer, Leverkusen, Germany) or an overdose of isoflurane (Abbot, Wiesbaden, Germany) followed by cervical dislocation.

### *Ex vivo* stimulation of tracheal tuft cells

Tracheal tuft cells were stimulated *ex vivo* as described previously^3^. More details can be found in the supplementary methods.

### Measurement of CysLTs and LTB_4_

CysLTs and LTB_4_ were measured in the tracheal supernatants and BALF using the commercially available Cysteinyl Leukotriene ELISA Kit (Cayman Chemical, Catalog no. Cay500390-96S) and the LTB_4_ Parameter Assay Kit (R & D Systems, Catalog no. KGE006B) according to the manufacturer’s protocol. A brief description can be found in the supplementary methods.

### Bacterial growth and preparation of bacterial supernatants

A detailed description of bacterial strains used in this study can be found in the supplementary methods^71^. Collection of bacterial supernatants was performed as previously indicated^2^ with a detailed description in the supplementary methods.

### Intratracheal administration of substances

The *in vivo* intratracheal administration of substances was performed as described previously^2^. A more detailed description can be found in the supplementary methods.

### Depletion of TRPV1 neurons

Ablation of TRPV1+ sensory neurons with resiniferatoxin (RTX, biomol, Hamburg, Germany) was performed as described previously^12^. A more detailed description is provided in the supplementary methods.

### Tissue preparation

The tissues were prepared as described previously^2^ and in the supplementary methods.

### Immunohistochemistry

Immunohistochemistry was performed on tracheal sections as previously indicated^2^. A detailed description of the procedures and antibodies used in this study is supplied in the supplementary methods.

### Analysis of Evans blue extravasation

The Evans blue extravasation was analyzed as described previously^2^. A detailed description can be found in the supplementary methods.

### Neutrophil migration assay

A detailed protocol of the neutrophil migration assay can be found in the supplementary materials.

### Measurement of CGRP

The estimation of CGRP in tracheal homogenates was carried out using the CGRP ELISA kit (Bertin Pharma, Catalog no. A05481) as indicated previously^2^. The supplementary method provides a detailed explanation of the procedures used.

### Infection of mice with *P. aeruginosa*

*Trpm5^+/+^* and *Trpm5^-/-^* mice were infected with the mucoid *P. aeruginosa* strain NH57388A (kindly provided by Niels Hoiby, Department of Clinical Microbiology, Rigshospitalet, University of Copenhagen, Denmark) as described previously^2,3^. A detailed description is provided in the supplementary methods.

### Single cell preparations and FACS analyses

Tracheae, BALF, blood, and spleen of *P. aeruginosa-infected* mice, and BALF obtained from mice treated intratracheally with bacterial supernatants, were collected and processed for FACS analyses as described previously^3^. A detailed description of the preparation and analyses, and gating strategies used can be found in the supplementary methods.

### Detection of bacterial eATP

ATP in bacterial supernatants was determined using the BacTiter-Glo™ Microbial Cell Viability Assay Reagent (Promega, Catalog no. G8230as), as described in the supplementary methods.

### LC-ESI-MS/MS measurement of bacterial quorum-sensing molecules

The measurements of bacterial QSMs using LC-ESI-MS/MS were performed as described previously^72^. A detailed description of the procedures used, together with parameters used for monitoring homoserine lactones (HSL), can be found in the supplementary methods.

### Bacterial killing assay

Bacterial survival of the *P. aeruginosa* strain NH57388A added to macrophages treated with tracheal supernatants was assessed as described previously^73^. A detailed description of the procedure used can be found in the supplementary methods.

### qRT-PCR

Quantitative RT-PCR was performed as previously described^3^. A brief description including primers used in this study is provided in the supplementary methods.

### Measurements of H_2_O_2_

A description of the H_2_O_2_ measurements using the AmplexTM Red Hydrogen Peroxide/Peroxidase Assay Kit (Invitrogen, Catalog no. A22188) can be found in the supplementary methods.

### MALDI-TOF mass spectrometry for analysis of bacterial pathogens

Bacterial swabs from mice were collected and subjected to analysis by MALDI-TOF. A detailed description of the procedure used can be found in the supplementary methods.

### Analysis of the single-cell RNA sequencing data

The previously published expression data (dataset GSE116525) for three sets of cell populations isolated from the murine tracheal epithelium were obtained from the Gene Expression Omnibus (https://www.ncbi.nlm.nih.gov/geo/). The data were *in silico* reanalyzed using the iDEP.96 as indicated previously^74^..

### Declaration of Generative AI and AI-assisted Technologies in the Writing Process

A Large Language Model (LLM), ChatGPT (GPT-5, OpenAI, San Francisco, CA, 2025), was used to improve grammar, readability, and style of the manuscript. All scientific content, interpretation, and conclusions were generated and verified by the authors, who take full responsibility for the final content of the manuscript.

### Statistical analyses

Each experiment included at least three biological and two technical replicates and was repeated at least three times. Multiple sections or samples per animal from a minimum of three animals were included for histological evaluation. The exact sample numbers and statistical tests are specified in the respective Figure legends. For the quantification of neutrophils in tissue sections, the cells were counted manually based on their fluorescent labeling using a fluorescence microscope. For the tracheae, five sections covering a distance of 5 mm (100 µm distance between each section) were evaluated. Statistical analyses were performed with GraphPad Prism 10.4.1 (GraphPad Software Inc., La Jolla, CA, USA). Data are depicted as means ± standard error of the mean with single values and an n-number of evaluated animals, samples, or cells. The Shapiro-Wilk test and the Kolmogorov-Smirnov test were used to assess the normal distribution of the data. Normally distributed data sets with more than two groups were then analyzed for significant differences using a one-way ANOVA, and non-normally distributed data with a Kruskal-Wallis test, both followed by Dunn’s post-hoc analyses for multiple comparisons. Direct comparisons between two groups were performed using the unpaired Student’s t-test for normally distributed data and with the Mann-Whitney test for non-normally distributed data (α = 0.05). Paired samples were analyzed with the paired Student’s t-test for normally distributed data and with the Wilcoxon test for non-normally distributed data. All data points represent measurements from distinct samples. The level of significance was set at p ≤ 0.05.

## ACKNOWLEDGMENTS

The authors would like to thank Andrea Rabung, Aline Herges, and Noreldin Mahmoud (Institute of Anatomy and Cell Biology, Saarland University, Homburg) for technical assistance. The authors would also like to thank Prof. Dr. Markus Hoth (Center for Integrative Physiology and Molecular Medicine (CIPMM), School of Medicine, Saarland University, Homburg, Germany, PD Dr. Elmar Krause (Department of Cellular Neurophysiology, Center for Integrative Physiology and Molecular Medicine (CIPMM), Saarland University, Homburg, Germany) for providing access to the FACS equipment, and Prof. Dr. Lorenz Thurner (Department of Oncology and Hematology, School of Medicine, Saarland University, Homburg, Germany) for providing using the Cytospin centrifuge. We would like to additionally thank Prof. Dr. Niels Hoiby (Department of Clinical Microbiology, Rigshospitalet, University of Copenhagen, Denmark), Prof. Dr. Bastian Opitz (Department of Infectious Diseases, Respiratory Medicine and Critical Care, Charité – Universitätsmedizin Berlin, Berlin, Germany), and Dr. Birgitt Gutbier (Department of Infectious Diseases and Pulmonary Medicine, Charité-Universitätsmedizin Berlin, Berlin, Germany) for providing us with bacterial strains used in the study. Creation of Figure. 1B, 2G, 4E and 8F was accomplished with the software Inkscape version 0.92.3. This study was supported by the German Research Foundation (DFG) SFB TRR152 grants P22 to GKC, P11 to UB, Z02 to UB, P14 to VC and TG, as well as by DFG KR 4338/1-2 to GKC, the Saarland University and Helmholtz-Institute for Pharmaceutical Research Saarland TANDEM initiative to AKHH, GKC, MB and ME, in addition to a HomFor2023 Nachwuchsförderung and HomForexzellent2023 grant from Saarland University to MIE.

## AUTHOR CONTRIBUTIONS

MIE, MG, NAW, PG, EH, GSL, FB, CK, AK, and NA performed experimental work. SK, AW, TG, VC, CS, and UB provided transgenic animals. SBE, MIH, AMK, SLB, MB, VF, ME, and AKHH provided resources and experimental tools. MIE, NAW, and GKC interpreted the data. MIE and GKC conceived and designed the study, interpreted the results, and wrote the manuscript. All authors have read the manuscript and approved its submission.

