## Supplemental data for "Tracheal tuft cell-released leukotrienes promote antibacterial immune responses"

**Supplementary Methods:**

#### *Ex vivo* stimulation of tracheal tuft cells

Cysteine leukotriene (CysLTs) and $LTB_4$ detection was performed using whole tracheae freshly explanted from euthanized mice. Tracheae were explanted and immediately incubated in a physiological buffer consisting of 5 mM KCl, 135 mM NaCl, 10 mM Glucose, 10 mM HEPES, 1 mM $MgCl_2$ and 2 mM $CaCl_2$. In some experiments, the buffer was supplemented with denatonium (in a final concentration of 1 mM) or ATP (in a final concentration of 10 mM) for stimulation of tuft cells. When indicated, the tracheae were incubated in the physiological buffer containing MK-886 (final concentration of 1 µM) or apyrase (5 U/ml) (Sigma-Aldrich) in addition to denatonium. The respective experimental conditions are indicated in the figures, the figure legends and the description of the results. After 10 minutes of incubation at 37°C, the supernatants were collected and used directly or stored at -80° C until further analysis.

#### Measurements of CysLTs and $LTB_4$

CysLTs were detected in the tracheal supernatants using the commercially available Cysteinyl Leukotriene ELISA Kit (Cayman Chemical, Catalog no. Cay500390-96S) according to the manufacturer's protocol. The detection strategy is based on a competitive reaction for binding CysLT ELISA Monoclonal Antibody between CysLTs and CysLTs-acetylcholinesterase conjugate. The lower limit of detection was 60 pg/ml. The following cross reactivity was reported: $LTC_4$ (100%), $LTD_4$ (100%), $LTE_4$ (79%), 5,6-DiHETE (3,7%), $LTB_4$ (1,3%), 5 (S)-HETE (0,04%) and AA ($<$0,01%). Briefly, samples, water or standard solution were added to the wells, followed by an addition of CysLT AChE Tracer to all wells except the blank and the total activity well. Within 15 minutes the CysLT ELISA monoclonal antibody was added everywhere, except to the blank, the total activity, and the non-specific binding wells. Then the plate was incubated overnight at 4 °C. The following day, the plate was washed 5 times with wash buffer, and 200 µl Ellman's Reagent was added to each well. Then a diluted tracer was added to the total activity well. The assay was developed in the dark on a horizontal shaker (VWR, Darmstadt, Germany) for 90 min and then read at 405 nm wavelength using a microplate reader (Bio-Rad, Laboratories GmbH, Feldkirchen, Germany).

For measuring $LTB_4$ in tracheal supernatants, the $LTB_4$ Parameter Assay Kit (R & D Systems, catalog no. KGE006B) was used. This assay is based on a competitive binding reaction of $LTB_4$ and a stable amount of horseradish peroxidase (HRP)-conjugated $LTB_4$ to bind with $LTB_4$ polyclonal antibody. The sensitivity of this reaction was 10.9 pg/ml, and no cross-species interactions were detected. Briefly, standard, control, sample, or buffer was added to the wells, and 50 µl of primary antibody solution was added to each well except the non-specific binding wells. The plate was incubated for 1 hour at room temperature on a horizontal shaker at approximately 500 rpm. Then, $LTB_4$ conjugate was added to each well, and the plate was incubated for 3 hours at room temperature on the shaker. The plate was washed 4 times with the wash buffer. Subsequently, substrate solution was added to each well, and the plate was incubated for 30 minutes at room temperature in the dark before adding the stop solution. Finally, the plate was read on a microplate reader (Bio-Rad, Laboratories GmbH, Feldkirchen, Germany) at 450 nm.

**Bacterial growth and preparation of bacterial supernatants**

Bacterial strains used in this study are listed in Supplementary Table 1. *P. aeuroginosa* strain NH57388A and *K. pneumonia* strain Kp52145 were cultivated on BD trypticase™ soy agar II with 5% sheep blood (BD) at 37 °C. *S. pneumoniae* strain PN36 was grown on Columbia agar supplemented with 5% sheep blood (BD

Biosciences, Heidelberg, Germany) for 8 h (37 °C, 5% CO2). Single colonies of *P. aeuroginosa* strain NH57388A and *K. pneumonia* strain Kp52145 were inoculated in LB broth (Luria/Miller) (Carl Roth) and grown overnight (12-16 h) under aerobic conditions, shaking at 225 rpm at 37 °C. For cultivating S. *pneumoniae* strain PN36, single colonies were inoculated in Todd-Hewitt broth (Oxoid, CM0189) supplemented with 0.5% yeast extract (BD Bioscience) and 10% heat-inactivated FBS (FBS Supreme, PAN-Biotech, Aidenbach, Germany). *S. aureus* strain Newman was grown in tryptic soy broth (TSB) (BD Biosciences). Main cultures were prepared by diluting the overnight cultures to an optical density at 600 nm ($OD_{600}$) of 0.05 and cultivating at 37 °C. The bacterial growth was monitored by measuring the $OD_{600}$ every two hours for a duration of 12-24 hours. Supernatants were collected by centrifugation at 16.000 rpm at 4 °C for 10 minutes. The collected supernatants were further filtered using 0.20 µM syringe filters (Sarstedt, Nümbrecht, Germany) to ensure bacterial removal, and they were used directly or kept frozen at -80 °C until use. For ATP depletion, bacterial supernatants were treated with apyrase (5 U/ml; Sigma-Aldrich) for 5 or 30 min at 37 °C.

**Intratracheal administration of substances**

Mice were narcotized with either urethane (1,5 gm/kg) (Sigma-Aldrich, Merck, Taufkirchen, Germany) or ketamine (90 mg/kg, Zoetis, Berlin, Germany)/xylazine (10 mg/kg, Bayer, Leverkusen, Germany) according to the respective animal protocol and were used for *in vivo* experiments to assess plasma extravasation and recruitment of neutrophils. Briefly, the anesthetized mice were injected with 100 µl Evans blue (4 mg/ml) into the systemic circulation via the retro-orbital venous sinus. Then, a midline incision was made in the ventromedial cervical region to expose the salivary glands, which were then positioned to a lateral position. To expose the trachea, the infrahyoid muscles were moved gently, and the median cricothyroid ligament of the larynx was identified. Four microliters of the following substances were applied directly into the tracheal lumen using a syringe (Hamilton, Reno, NV, USA): denatonium benzoate (1, 10, or 20 mM; Sigma-Aldrich) or bacterial supernatants of the *P. aeruginosa* strain NH57388A or the corresponding vehicle solutions (PBS; Gibco, Thermo Fisher Scientific, and LB broth; Carl Roth). When indicated, bacterial supernatants were applied simultaneously with 5 U/ml apyrase (Sigma-Aldrich). In experiments where the MK-886 was used, animals were injected with vehicle (PBS) or 10 mg/kg MK-886 (Merck Millipore) 30 minutes before the intratracheal injection of substances. All animals were euthanized 30 minutes after stimulation of the tracheal tuft cells using inhalation of an overdose of isoflurane (Piramal Critical Care Deutschland GmbH, Hallbergmoos, Germany) followed by cervical dislocation.

### Depletion of TRPV1 neurons

Resiniferatoxin (RTX) was used in three increasing doses (30, 70, and 100 µg/kg) applied on three successive days to deplete TRPV1-expressing neurons. One day before injecting the mice, Tramal (Grünenthal, Aachen, Germany) was given in the drinking water at a concentration of 1 mg/ml. The subcutaneous injections of RTX were performed under isoflurane narcosis (with an initial flow rate of 1 l/min at a concentration of 3%, followed by 2% for 3 minutes). The RTX-treated mice were used for intratracheal application of substances as described above one week after the last RTX injection.

### Tissue preparation

Mice were euthanized using inhalation of an overdose of isoflurane (Piramal Critical Care Deutschland GmbH, Hallbergmoos, Germany) followed by cervical dislocation.

Following euthanization, animals were transcardially perfused with Zamboni fixation solution. Tracheae were explanted and divided into three segments (1-5, 6-11 tracheal cartilaginous rings, and 12-bifurcation). Then, tracheae were postfixed with Zamboni, washed, and incubated with 18% sucrose before being shock frozen. 10 µm thick cryosections distanced 100 µm from each other were prepared from the intermediate tracheal segment (6-11 tracheal cartilaginous rings) and used to analyze Evans blue extravasation and neutrophil recruitment.

**Immunohistochemistry**

Briefly, tracheal sections were blocked for 1 hour and incubated overnight at room temperature with the following primary antibodies: anti-Ly6G-FITC antiserum (Thermo Fisher Scientific, 11-5931-85, RRID: AB_465315; 1:200) or rabbit-anti-Trpm5-antiserum (generated in-house and validated in tissue from Trpm5-deficient mice[1]. The next day, the tissue sections were washed three times in PBS and incubated for 1 hour with a donkey anti-rabbit antibody coupled to Cy3 (Merck Millipore, AP182C, Lot 2548999; 1:1000) at room temperature. The sections were incubated for 10 minutes with DAPI, followed by three washing steps with PBS, and then mounted with Mowiol. Evaluation of the infiltrating neutrophils as well as the Trpm5$^{+}$ tuft cells was performed by counting the cells per tracheal ring manually using an epifluorescence microscope (Axioplan 2 imaging, Zeiss, Jena, Germany).

**Analysis of Evans blue extravasation**

The extravasation of Evans blue within the tracheal cryosections was evaluated using an epifluorescence microscope (Axioplan 2 imaging, Zeiss) by using a Texas Red filter (585/29 nm excitation; 624/40 nm emission). Images of the whole tracheal rings were obtained using the same exposure time for all experimental conditions. The Evans blue intensity in the lamina propria was assessed by measuring the fluorescence intensity of each ring using the ImageJ software (NIH).

**Neutrophil migration assay**

The previously isolated mouse lung endothelial cells (see above, FACS sorting) were seeded at a density of 1 x $10^4$ in 100 µl on the top of 6.5 mm Transwell filters in 24-well plates (Corning permeable support, 0.4 µM polyester membrane). Cells were cultivated in complete medium consisting of DMEM (Gibco, Thermo-Fisher), 20 % heat inactivated FBS (FBS Supreme, PAN-Biotech, Aidenbach, Germany), 1X Endothelial cells growth supplement (ECGS; Sigma-Aldrich), 100 U/ml penicillin and 100 mg/ml streptomycin (Sigma-Aldrich). The transwell inserts were incubated at 37˚C overnight under liquid/liquid conditions with 600 µl of DMEM supplemented with 10% (v/v) heat-inactivated FBS (FBS Supreme, PAN-Biotech, Aidenbach, Germany) and 1% (v/v) Antibiotic-Antimycotic (Gibco, Thermo Fisher Scientific) in the basolateral compartment and 50 µl of the same media in the apical compartment. The monolayer formation was confirmed by eye using an upright light microscope. The integrity and permeability of the monolayer were additionally estimated by measuring the transepithelial electrical resistance (TEER) using an EVOM2 device (WPI, Friedberg, Germany). Bone marrow cells were isolated from euthanized wild type mice by flushing the bone marrow out of the femur and tibia with PBS. RBCs were lysed by adding 1 ml ACK lysis buffer (Gibco) for 15 minutes before adding 10 ml PBS (Gibco, Thermo Fisher Scientific). Bone marrow cells were centrifuged at 250 x *g* at room temperature for 5 minutes. Sedimented bone marrow cells were resuspended into 1 ml DMEM, and cell counts were determined using NucleoCounter® NC-200 (ChemoMetec). Differential cell staining with performed using RAL Diff-Quik™ Kit (Cellavision, Lund, Sweden) according to the manufacturer's recommendations. 100 µl of the resuspended bone marrow cells were applied on the proliferated endothelial cell layer and allowed to migrate for two hours toward tracheal supernatants obtained from 1 mM denatonium-stimulated tracheae or agonists including CGRP (Sigma) and $LTB_4$ (Sigma). Calculation of the percentages of migrated neutrophils was carried out using differential cell staining as described above counting 200 cells.

**Measurement of CGRP**

Mice were euthanized with isoflurane followed by cervical dislocation. Tracheae were explanted and collected in 200 µl physiological buffer using 2 ml Eppendorf tubes. Stimulation of explanted tracheae was performed with denatonium for 5 minutes at 37°C before being homogenised on ice using a tissue ruptor (Qiagen, Hilden, Germany) and centrifuged at 1500 g for 5 min at room temperature. Supernatants were collected and transferred into new 1.5 ml Eppendorf tubes and processed directly for measuring CGRP using CGRP ELISA Kit (Bertin Pharma, Montigny-leBretonneux, France) according to the manufacturers' protocols. Plates were assessed photometrically using a microplate reader (Bio-Rad, Laboratories GmbH, Feldkirchen, Germany).

**Mice infection with *P. aeruginosa***

Briefly, mice were anesthetized with ketamine/xylazine (90-120 mg/kg / 6-8 mg/kg; Zoetis/Bayer) and 40 µl of bacterial suspension of *P. aeruginosa* strain NH57388A ($10^6$ CFU) were applied intratracheally. To deplete LTs, mice were injected with MK-886 (10 mg/kg). Mice were sacrificed with an overdose of ketamine/xylazine (five times higher than the dose used for anesthesia after the mice were anesthetized with a single dose) 4 hours after the inoculation and organs were collected for FACS analyses.

**Single cell preparations and FACS sorting**

Briefly, Blood was first collected using EDTA coated canula from the vena cava and stored at 4°C till further processing. Analysis of the blood composition was performed by FACS or be staining blood smears using Diff-Quik™ Kit (Cellavision, Lund, Sweden). The BALF was then collected by rinsing the lungs of euthanized mice three times with 1 ml of ice cold FACS buffer (0.1 mM EDTA and 1% FBS in DPBS) per mouse through an intratracheal canula. The mice were then perfused with 10 ml of DPBS per mouse through the left ventricle of the heart and the tracheae were dissected from the larynx till the level of the bifurcation. For preparation of single cell suspensions, the tracheas were digested first in dispase solution (15 U/ml) and DNaseI (150 U/ml) for 30 minutes at room temperature. Following that, the tracheae were cut into small pieces and further digested in collagenase I solution (1 mg/ml) and DNaseI (150 U/ml) for 30 minutes at 37 °C. The cells were then passed through a 70 µm cell strainer.  Spleens were also collected and mechanically disrupted and filtered through a 70 µm cell strainer to prepare a single cell suspension. Blood, trachea and spleen cells were treated with ACK lysis buffer (Thermo Fisher Scientific) to remove red blood cells. Cell counts and viability were measured using an automatic cell counter (NucleoCounter® NC-200™, Chemomatec, Kaiserslautern, Germany). The collected cells were fixed using 1% PFA and subsequently stained. For staining, Fc block (CD16/CD32 antibodies) were first added to the cells for 30-45 minutes followed by adding fluorophore conjugated antibodies cocktails for 45 minutes at 4 °C. The antibodies used in the study are listed in Supplementary Table 2 The FACS measurements were performed and analyzed using a BD FACSverse machine (BD Biosciences, Heidelberg, Germany) and BD FACSuite™ Software version 1.0.6.5230 (BD Biosciences). Data analysis was achieved using FlowJo v10.9.0 (FlowJo, LLC, BD Biosciences).

For isolation of lung endothelial cells, wild type mice were euthanized, and the lungs were perfused through the right ventricle of the heart with ice cold DPBS. The lungs were dissected and cut into small pieces and digested in 4 ml of collagenase I (1 mg/ml) and DNaseI (150 U/ml) for 45 minutes. Afterwards, the cells were filtered with a 70 µm cell strainer and the RBCs were removed using ACK lysis buffer. Fc block (CD16/CD32 antibodies) was first added to the cells followed by addition of anti CD31 and CD45 (PerCp-Cy5.5) for 30 minutes at 4 °C. Then an Alexa Fluor 488 secondary antibody (Invitrogen) was added for 30 minutes. The cell sorting was performed on the FACS Aria III (BD Biosciences).

**Detection of ATP**

Extracellular ATP released in bacterial supernatants was determined using the BacTiter-Glo™ Microbial Cell Viability Assay Reagent (Promega, Walldorf, Germany, catalog no. G8230) following the manufacturer's guidelines and as described previously (Mempin et al., 2013). Briefly, 100 µl of filtered bacterial supernatants were mixed with an equal volume of BacTiter-Glo™ Microbial Cell Viability Assay Reagent in a 96 well plate and incubated at room temperature for 5 minutes before measuring the luminescence at a wavelength of 600 nm using the EnSight Multimode Plate Reader (Perkin Elmer™, Rodgau-Jügesheim, Germany) with the software Kaleido 2.0 (Perkin Elmer).

**LC-ESI-MS/MS measurement of bacterial quorum-sensing molecules**

Supernatants obtained at different time points (0, 2, 4, 6, 8, 10, and 12 h) were collected from bacterial cultures grown in LB medium at 37 °C with shaking at 225 rpm. Alkylquinolones and homoserine lactones (HSL) were quantified via LC-ESI-MS/MS analysis using a Dionex Ultimate 3000 HPLC system coupled to a TSQ Quantum

Access MAX (Thermo Fisher Scientific, Waltham MA). Alkylquinolone quantification was performed as described previously (Hamed et al., 2023) using the mass transitions as given in Supplementary Table 3. For C4-HSL, C6-HSL was used as an internal standard, and the following conditions were applied: Luna Omega 3 µM Polar C18 100 Å, 150 x 2.1 mm (Phenomenex, Torrance, CA, USA); eluent A – $H_2O$ with 0.1% trifluoroacetic acid (TFA), 0.1% pentafluoropropionic acid (PFPA), 0.1% heptafluorobutyric acid (HFBA); eluent B – acetonitrile with 0.1% TFA, 0.1% PFPA, 0.1% HFBA; flow 0.5 mL/min. 0 – 0.9 min 5% B; 0.9 – 2 min 5% B – 70%; 2 – 4 min 70% B – 90% B; 4 – 5.5. min 90% B; 5.5 – 6 min 5% B. MS instrument parameters: spray voltage 4500 V, vaporizer temperature 297 °C, sheath gas pressure 20 psi, aux gas pressure 55 psi, capillary temperature 270 °C, collision pressure 1.5 mTorr, positive ionization mode. For HSL, analyte reaction monitoring parameters are shown in Supplementary Table 3. Data acquisition and quantification were performed using Xcalibur software 4.2 with the use of a calibration curve relative to the area of the internal standard.

**Bacterial killing assay**

To assess the bacterial survival capacity upon being ingested by lung interstitial macrophages, we used a tetrazolium reduction assay as indicated previously (Ofek et al, 2011). Briefly, macrophages were plated at a density of 1 x $10^6$ cells/ml in 96 well plates. The next day, cells were treated with tracheal supernatants obtained from *Trpm5*$^{+/+}$, *Trpm5*$^{-/-}$ and *Trpm5*-DREADD mice which were stimulated with 1 mM denatonium or 100 µM CNO, respectively. Macrophages were stimulated with tracheal supernatants for 30 minutes before being infected with the *P. aeruginosa* strain NH57388A at a multiplicity of infection (MOI) of 20:1. After 60 minutes, the extracellular bacterial cells were removed by washing twice with PBS (Gibco, Thermo Fischer

Scientific), and macrophages were lysed by adding 0.1% Triton X-100 (Carl Roth). The bacterial survival was assessed by measuring the optical density (OD595) after adding 3-(4,5-Dimethyl-2-thiazolyl)-2,5-Diphenyl-2H-Tetrazolium-Bromid (MTT) (Merck) in a concentration of 5 mg/ml for 30 minutes. Assessment of the number of intracellular bacterial cells was performed by measuring the absorbance intensity at a wavelength of 595 nm using the EnSight Multimode Plate Reader (Perkin Elmer™, Rodgau-Jügesheim, Germany) with integrated software Kaleido 2.0 (Perkin Elmer). The measured intensity correlated with the counts of macrophage-associated intracellular bacterial cells. Survival results were expressed as a percentage of the ingested bacterial cells, where the percentages correspond to 100% X A595 control plate/experimental plate.

**qRT-PCR**

Isolation of RNA of FACS-sorted cells was performed using the RNAqueousTM-Micro Kit (Invitrogen; catalog no. AM1931) according to the manufacturer's protocol. The genomic DNA (gDNA) was digested using the DNase reagents provided with the kit. Isolation of RNA from whole tracheal preparations was performed using the RNeasy Mini Kit (Qiagen, Hilden, Germany; catalog no. 74104) according to the manufacturer's protocol. The concentration and purity of the isolated RNA was evaluated using a NanoDrop instrument (Thermo Scientific, Waltham, MA, USA). Elimination of contaminating gDNA within the RNA samples isolated from whole tracheal preparations was performed as follows: 1 µg total RNA, or 8 µl in samples with low concentration, were incubated with 1 µl DNaseI (Thermo Scientific) within 1 µl 10x DNaseI buffer (Thermo Scientific) for 30 minutes at 37 °C. The reaction was stopped by adding 1 µl 25 mM EDTA (Thermo Scientific) and incubated at 65 °C for 10 minutes. The subsequent cDNA synthesis was performed using the SuperScript II reverse transcription kit (Thermo Scientific, REF: 18064014) according to the instructions of the manufacturer. Briefly, 600 µg DNase-digested total RNA or 10 µl in cases of lower concentration, were mixed with 1 µl Oligo $dT_{18}$ Primer (Thermo Scientific) and 1 µl dNTPs (Thermo Scientific). Reactions were incubated for 5 minutes at 65°C. 4 µl First strand Buffer (Thermo Scientific) and 2 µl DTT (0.1 M; Thermo Scientific) were added to the reactions and incubated for 2 min at 42 °C. Reverse transcription was initiated by adding 1 µl of SuperScript II Reverse Transcriptase. Reactions without enzyme were run in parallel serving as controls. Reactions were run for 50 minutes at 42 °C, before being terminated at 72 °C for 15 minutes.

Gene expression was quantified using the SYBR®Green detection method using specific primers listed in Supplementary Table 4. The qRT-PCR reactions were performed using a Bio-Rad CFX Connect™ RealTime System (Bio-Rad Laboratories, Inc., Hercules, CA, USA) and consisted of 10 µl iTaq Universal SYBR®Green Supermix (Bio-Rad), 6.6 µl nuclease-free water, 2.4 µl specific primer mix (with a final concentration of 10 mM for each primer) and 1 µl cDNA (1 ng/µl). Samples without reverse transcriptase were used as negative controls. The reactions were incubated at 95 °C for 30 s followed by a repetition of 40 cycles of 5 s at 95 °C and 30 s at 60 °C. PCR products were visualized by being separated on a 2-2.5% agarose gel and photographed using a ChemiDoc™ XRS (Bio-Rad). The relative expression analysis was performed using the $2^{-(\Delta\Delta CT)}$ method with the $\Delta$CT = CT (gene of interest; GOI) - CT (housekeeping gene), where the housekeeping genes used were Glyceraldehyde-3-phosphate dehydrogenase (*Gapdh*) for measuring the relative expression of genes in RNA samples isolated from macrophages and beta-2-microglobulin (*B2m*) for all other RNA samples.

**Measurements of $H_2O_2$**

Lung interstitial macrophages were plated in 96-well plates in the same density described for the bacterial killing assay. Macrophages were treated with tracheal supernatants obtained from wild-type or Trpm5-deficient mice under different conditions. The generated $H_2O_2$ was measured using the Amplex™ Red Hydrogen Peroxide/Peroxidase Assay Kit (Invitrogen, catalog no. A22188) according to the instructions of the manufacturer.

**MALDI-TOF mass spectrometry for analysis of bacterial pathogens**

Bacteria were collected in sterile PBS (Gibco, Thermo Fisher Scientific) using sterile cotton swabs inserted into the trachea of euthanized mice. Bacteria were identified using a MALDI-TOF target plate (Bruker Daltonics, Billerica, MA, USA). Briefly, bacterial suspensions were spotted on the plate. The dried spots were overlaid with matrix solutions. A 1 µl α-cyano-4-hydroxycinnamic acid (CHCA) matrix solution (Bruker Daltonics) comprising 50% (v/v) saturated CHCA, 2.5% (v/v) trifluoroacetic acid, and 47.5% (v/v) LC-MS grade water was applied on top of the dried spots. Bacterial Test Standard (BTS) (Bruker) was used for the calibration. Measurement was done using a Microflex LT Mass Spectrometer (Bruker Daltonics). To create the mass profiles in linear positive ion mode, 240 laser pulses were used at random locations in six different spots in the target plate. For the measurement, mass charge ratio ranges (m/z) between 2 KDa and 20kDa were created using a laser frequency of 60 Hz and a high voltage of 20 kV. After eliminating all flatlines and outlier peaks, the raw spectra were acquired and examined using Bruker Daltonics' FlexAnalysis® software version 3.4.

**Supplementary Table 1: Table of bacterial strains used in this study**

| Strain | Description | Reference or Source |
|---|---|---|
| *P. aeuroginosa* NH57388A | mucoid cystic fibrosis clinical isolate | Kindly provided by Prof. Niels Hoiby, Department of Clinical Microbiology, Rigshospitalet, University of Copenhagen, Denmark |
| *K. pneumoniae* strain Kp52145 | Reference strain; sequence type ST66, capsular serotype K2. | Kindly provided by Prof. Dr. Bastian Opitz, Department of Infectious Diseases, Respiratory Medicine and Critical Care, Charité-Universitätsmedizin Berlin, Berlin, Germany and German Center for Lung Research (DZL), Berlin, Germany |
| *S. pneumoniae* PN36 | Reference strain; NCTC 7978, serotype 3 | Kindly provided by Dr. Birgitt Gutbier, Department of Infectious Diseases and Pulmonary Medicine, Charité-Universitätsmedizin Berlin, Berlin, Germany |
| *S. aureus* Newman | ATCC 25904, wildtype | Gannoun-Zaki et al., 2018 |

**Supplementary Table 2: Table of antibodies used for immunohistochemistry and FACS**

| Antibody name | Dilution | Species/Clone | Supplier/Source | RRID or Cat. No. |
|---|---|---|---|---|
| TRPM5 | 1:400 | rabbit/polyclonal | VF | generated in house |
| TRPV1 | 1:1600 | rabbit/polyclonal | Alomone labs | ACC-030 Lot no. ACC030AN3525 |
| CD31 | 1:400 | rat/monoclonal | dianova | DIA-310 Lot 15916/05 |
| LY6G-FITC | 1:200 | rat/monoclonal | Life technologies | RM3001 Lot 1647833 |
| donkey antirabbit Cy3 | 1:1000 | donkey | Merck Millipore | AP182C Lot 2548999 |
| donkey antirabbit Alexa Fluor 488 | 1:500 | donkey | invtrogen | A48269 Lot ZG395855 |
| CD16/32 unconjugated | 1:100 | 93 | BioLegend | AB_312801 |
| Ep-CAM-PE-Cy7 | 1:50 | G8.8 | Biolegend | AB_1236477 |
| F4/80-PE-Cy7 | 1:40 | BM8 | BioLegend | AB_893478 |
| CD45-PerCPCy5.5 | 1:133 | 30-F11 | BioLegend | AB_893340 |
| CD11b-FITC | 1:40 | M1/70 | BioLegend | AB_312789 |
| CD86-PE | 1:133 | GL1 | BioLegend | AB_313151 |
| CD11c-APC-Cy7 | 1:40 | HL3 | BD Bioscience | AB_10611727 |
| LY6G-APC | 1:40 | 1A8 | BioLegend | AB_2227348 |
| CD45-APC-Cy7 | 1:40 | 30-F11 | BioLegend | AB_312981 |
| F4/80-FITC | 1:66 | BM8 | BioLegend | AB_893502 |
| CD11b-Pacific Blue | 1:50 | M1/70 | BioLegend | AB_755985 |
| MHCII-PE-Cy7 | 1:200 | N5/114.15.2 | BioLegend | AB_2069376 |
| CD206-PerCPCy5.5 | 1:40 | C068C2 | BioLegend | AB_2561992 |
| CD80-brilliant violet510 | 1:50 | 16-10A1 | BioLegend | AB_2810337 |
| CD121a (IL-1R)-APC | 1:20 | JAMA-147 | BioLegend | AB_2264757 |

**Supplementary Table 3: LC-ESI-MS/MS reaction monitoring parameters of HSL analytes in positive ionization mode**

| Molecule | Precursor ion m/z | Product ion m/z | Collision energy (V) | Tube lens offset (V) |
|---|---|---|---|---|
| PQS | 260.048 | 145.958 | 44 | 110 |
| | | 174.927 | 30 | 110 |
| HHQ | 244.050 | 158.944 | 31 | 100 |
| | | 171.943 | 33 | 100 |
| HQNO | 260.036 | 130.089 | 45 | 111 |
| | | 158.908 | 28 | 111 |
| 2-AA | 136.016 | 91.048 | 24 | 68 |
| | | 117.998 | 13 | 68 |
| DHQ | 161.971 | 115.979 | 27 | 84 |
| | | 143.941 | 19 | 84 |
| d4-HHQ | 248.081 | 162.965 | 32 | 100 |
| | | 175.982 | 34 | 100 |
| C4-HSL | 172.097 | 71.200 | 13 | 87 |
| | | 102.200 | 10 | 87 |
| C6-HSL | 200.118 | 99.200 | 11 | 88 |
| | | 102.100 | 10 | 88 |

**Supplementary Table 4: Table of primers used for qRT-PCR**

| Primer | Direction | Sequence (5’-3’) | Nucleotide reference |
|---|---|---|---|
| *Trpm5* | For. | TGAGGAACGACCTTTGGCTA | NM_020277.2 |
| *Trpm5* | Rev. | ACACGGATCTTGGTGGATGT | |
| *Pla2g4a* | For. | CAGCCACAACCCTCTCTTACTTC | NM_008869.4 |
| *Pla2g4a* | Rev. | CGGCATTGACCTTTTCCTTC | |
| *Alox5* | For. | TGTTCCCATTGCCATCCAG | NM_009662.2 |
| *Alox5* | Rev. | CACCTCAGACACCAGATGCG | |
| *Alox5ap* | For. | TGTTCCTCATGTCCTTCGCC | NM_009663.2 |
| *Alox5ap* | Rev. | GAGATCGTCGTGCTTACCGT | |
| *Ltc4s* | For. | CTCTTCTGGCTACCGTCACC | NM_001313968.2 |
| *Ltc4s* | Rev. | AAGCCCTTCGTGCAGAGAT | |
| *Lt4ah* | For. | AAGGGTCACCGATGGAAATC | NM_008517.2 |
| *Lt4ah* | Rev. | CTGGCACTGACTGAAGAGATAC | |
| *Cysltr1* | For. | TGAGGTACCAGATAGAGGCTG | NM_001281859.1 |
| *Cysltr1* | Rev. | CTTGGTGCCTTGGAGGTACA | |
| *Cysltr2* | For. | GGTGGATATGGGCAAGTAACA | NM_001162412.2 |
| *Cysltr2* | Rev. | GGTGACAGACCACCTTCTAATC | |
| *Ltbr1* | For. | GATGGCTGCAAACACTACATC | NM_008519.3 |
| *Ltbr1* | Rev. | CAGGATGCTCCACACTACAA | |
| *Ltbr2* | For. | CTTGACTCATCCTCTCCGCA | NM_020490.3 |

| | | | |
|---|---|---|---|
| *Ltbr2* | Rev. | TTCTAGGGGGAGAGCCGC | |
| *Nox2* | For | CGGAGAGTTTGGAAGAGCATAA | NM_007807.5 |
| *Nox2* | Rev | GGTACTGGGCACTCCTTTATTT | |
| *B2m* | For. | ATTCACCCCCACTGAGACTG | NM_009735.3 |
| *B2m* | Rev. | GCTATTTCTTTCTGCGTGCAT | |
| *Gapdh* | For. | AGGTCGGTGTGAACGGATTTG | NM_001411840.1 |
| *Gapdh* | Rev. | TGTAGACCATGTAGTTGAGGTCA | |

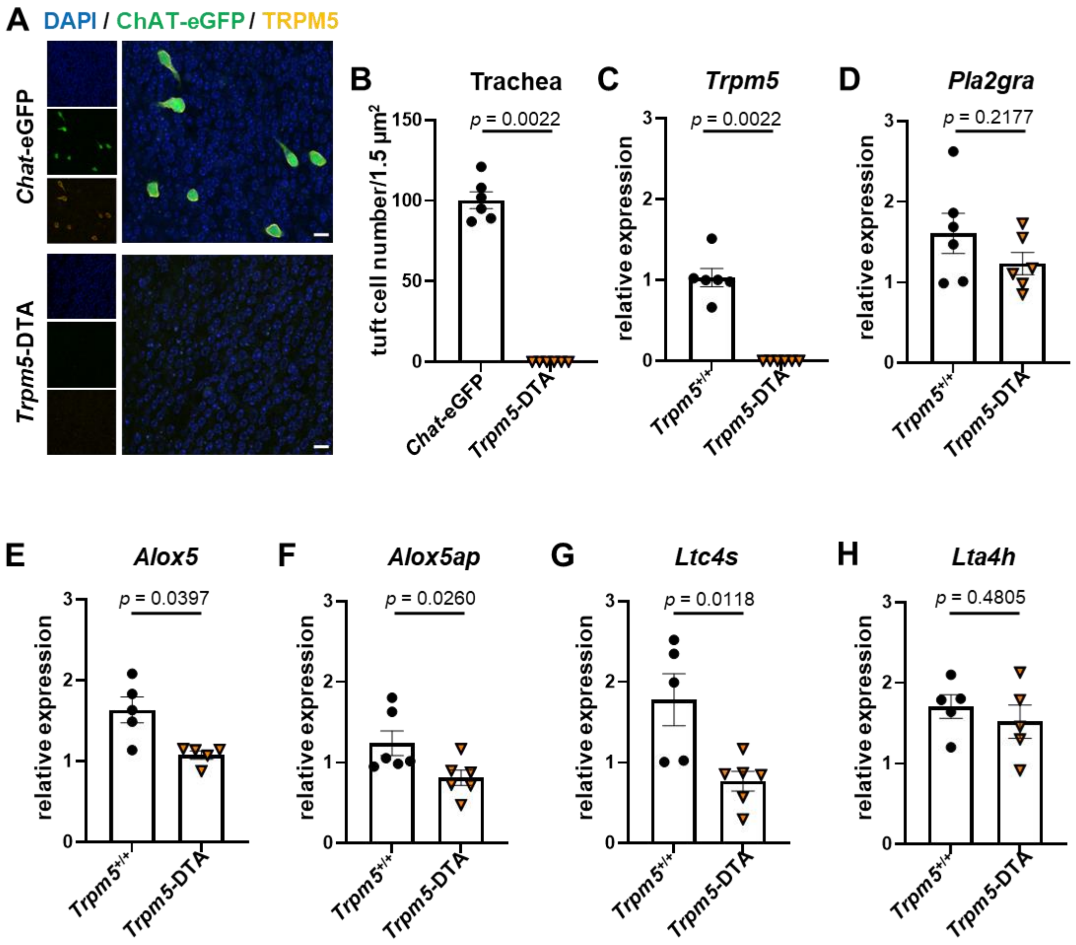


**Supplementary Figure 1: Tracheal tuft cells express genes required to synthesize LTs.** (A) Whole mount staining for Trpm5 (orange) of tracheae obtained from *Chat*-eGFP (green) and *Trpm5*-DTA mice. Scale bar, 20 µm. (B) Quantification of tracheal Trpm5$^{+}$ cells in entire whole mount stainings from *Chat*-eGFP (n=6) and *Trpm5*-DTA (n=6) mice. (C) Quantitative reverse transcription polymerase chain reaction (qRT-PCR) of whole tracheae for *Trpm5 (n=6)*. (D-H) Relative expression of *Pla2gra*, *Alox5*, *Alox5ap*, *Ltc4s,* and *Lta4h* in whole tracheae from *Trpm5*$^{+/+}$ (n=5-6) and *Trpm5*-DTA (n=5-6) mice. Data represent SEM of 5-6 independent experiments (each symbol represents a single animal). Two-tailed Mann-Whitney test (B, C, E, and F) or two-tailed unpaired Student's t-test (D, G, and H).

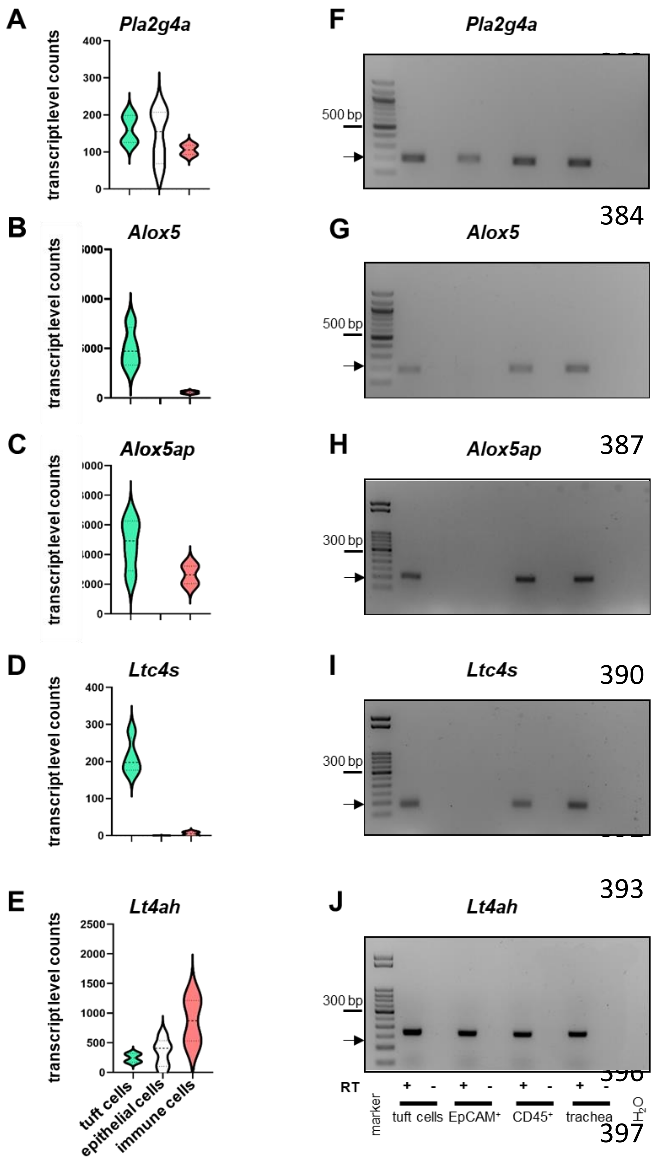


**Supplementary Figure 2: Tracheal TCs express genes required for the synthesis of LTs.** (A to E) Reanalysis of single RNA-sequencing data (dataset GSE116525) of *Pla2gra*, *Alox5*, *Alox5ap*, *Ltc4s,* and *Lta4h* in the three sets of cells (tuft cells, tracheal epithelial cells, and immune cells). (F-J) One representative gel image of five qRT-PCR experiments of LT-synthesizing genes in the same three populations of cells. Tracheal homogenates and $H_2O$ were used as controls. Additional samples were run without RT (reverse transcriptase).

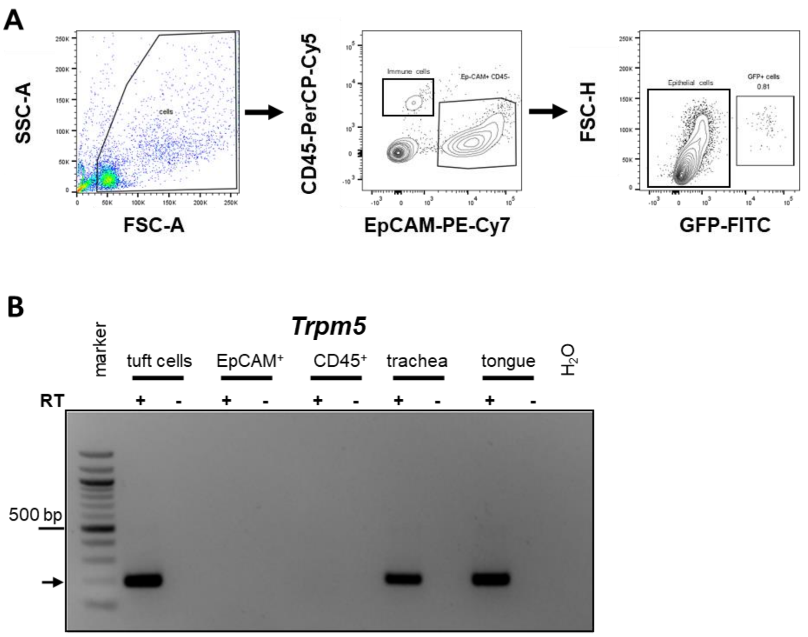


**Supplementary Figure 3: Trpm5 is exclusively expressed in tracheal TCs**. (A) Gating strategy for TCs (identified by eGFP expression within EpCAM$^+$ CD45$^-$ population), epithelial cells (EpCAM$^+$, CD45$^-$, GFP$^-$), and immune cells (EpCAM$^-$, CD45$^+$). (B) qRT-PCR for *Trpm5* of cells isolated by FACS-sorting, agarose gel electrophoresis for *Trpm5* transcript for tuft cells, epithelial cells, and immune cells. Whole tracheae and $H_2O$ served as positive and negative controls, respectively. The possibility of DNA-nonspecific binding of primers was excluded by running samples without RT (reverse transcriptase).

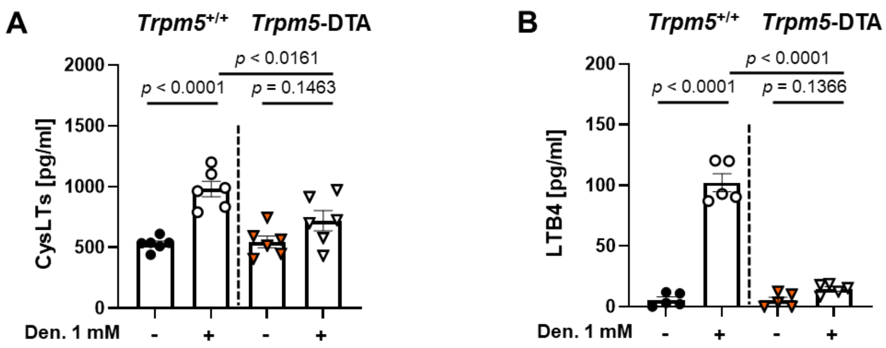


**Supplementary Figure 4: Tracheal tuft cells are the main epithelial source of CysLTs and LTB4 released upon tuft cell stimulation.** ELISA measurements of CysLTs (A) and LTB4 (B) in tracheal supernatants from *Trpm5*$^{+/+}$ and *Trpm5*-DTA mice after stimulation with 1 mM denatonium (Den.). Data represent mean ± SEM of 5-6 independent experiments per group. One-way ANOVA followed by Bonferroni's multiple-comparison test.

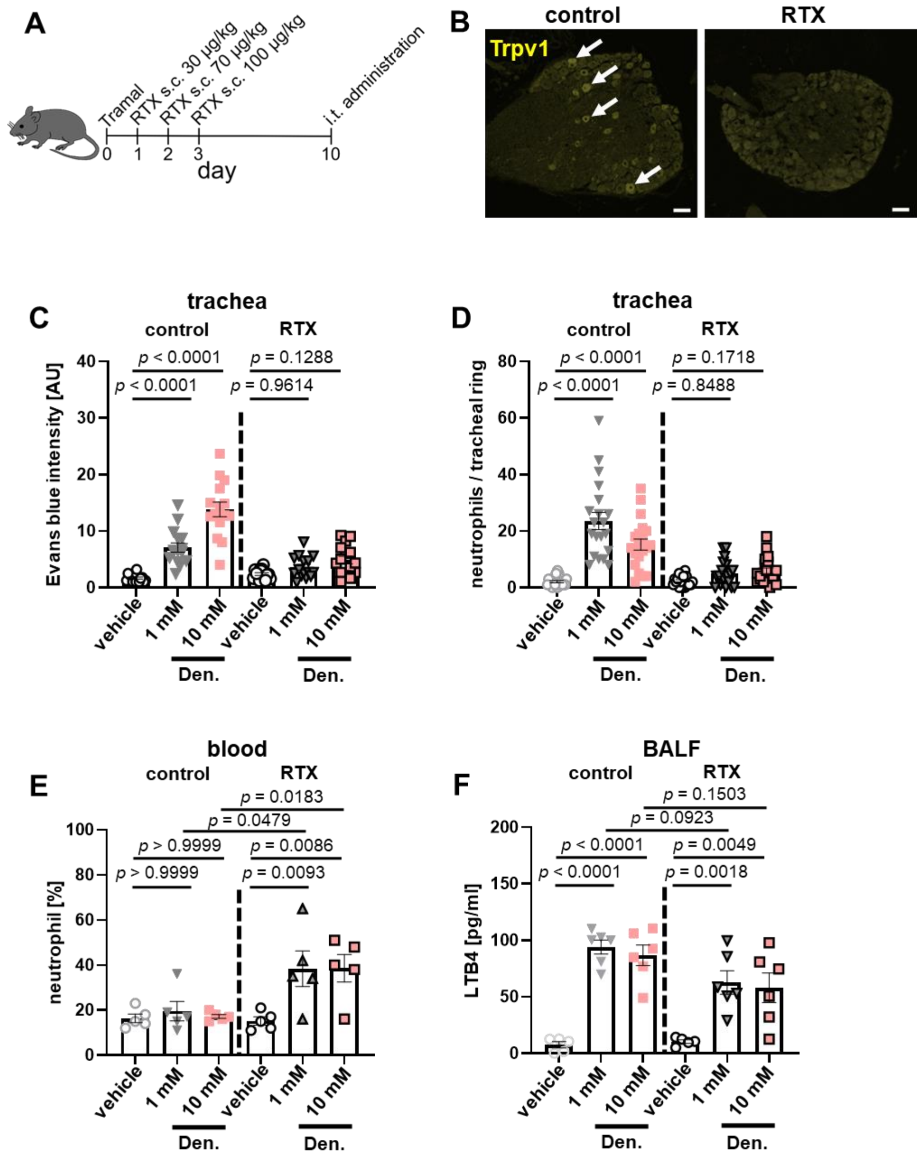


**Supplementary Figure 5: Tracheal tuft cells induce neurogenic inflammatory responses.** (A) Schematic of Resiniferatoxin (RTX) administration protocol for sensory neuron depletion in mice. (B) Representative immunohistochemical staining of Trpv1 in dorsal root ganglia (DRG) of control (PBS injected) and Resiniferatoxin (RTX)-injected mice. Scale bar, 50 µM. Arrows: Trpv1$^{+}$ neurons. (C) Quantification of Evans blue (EB) intensity in tracheal rings in response to vehicle (PBS), 1 and 10 mM denatonium (Den.) in control (n=4) and RTX-groups (n=4). (D) Quantification of neutrophils in tracheal rings of the same mice (E) Analysis of neutrophil percentages in the blood of control (n=5) and RTX-treated mice (n=5). (F) ELISA measurement of LTB4 in bronchoalveolar lavage fluids (BALFs) (n=5-6). Data represent means ± SEM of 15-20 tracheal rings of 4 mice (B-C) and 5-6 mice (D-F). Kruskal-Wallis with Dunn's test for pairwise multiple comparisons (B-C) and One-way ANOVA followed by Bonferroni's multiple-comparison test (E-F).

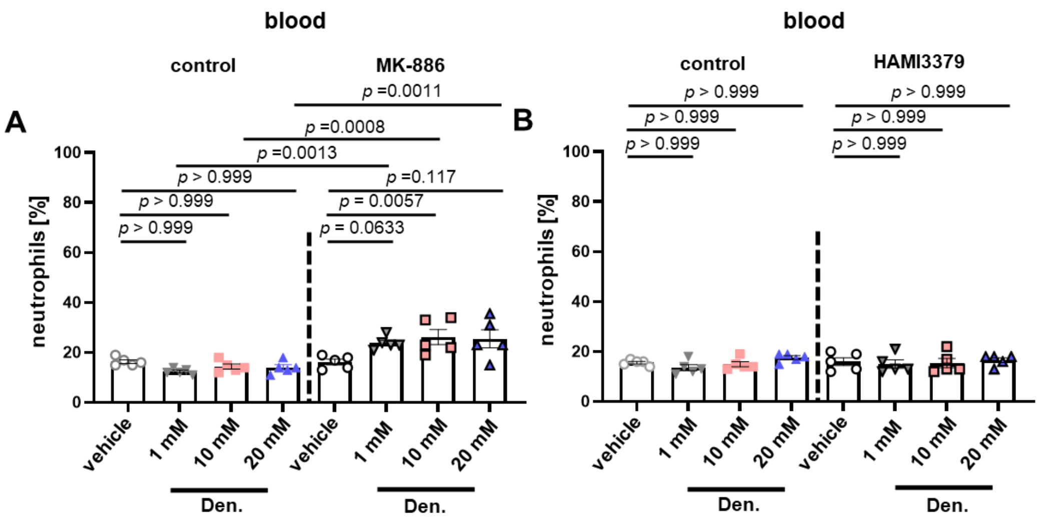


**Supplementary Figure 6: LTB4 released from tracheal tuft cells is required for the recruitment of neutrophils from the circulation.** (A and B) Analysis of blood neutrophils following the application of vehicle (PBS) and denatonium (Den.) in control groups (n=5) and mice treated with MK-886 (n=5) (A) or HAMI3379 (n=5) (B). Data are means ± SEM of 5 mice (A-B). Kruskal-Wallis with Dunn's test for pairwise multiple comparisons (A-B).

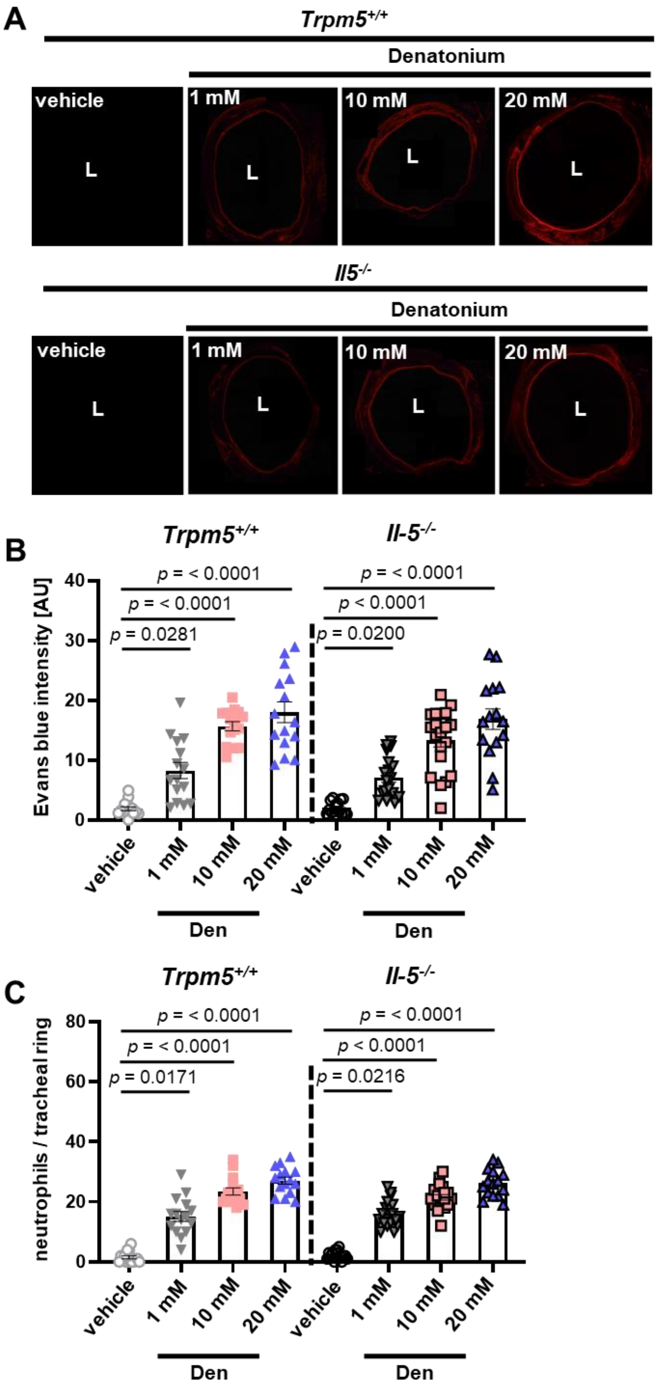


**Supplementary Figure 7: IL-5 signaling is not involved in the tuft cell-induced neurogenic inflammation.** (A) Representative image of EB extravasation in tracheal rings of wild type and *IL-5*$^{-/-}$ mice following the application of vehicle (PBS) and denatonium (Den.; 1, 10, and 20 mM). L, lumen. (B) Quantification of Evans blue extravasation in tracheal rings of wild type (n=4) and *IL-5*$^{-/-}$ (n=4) mice after tuft cell activation. (C) Analysis of neutrophils per tracheal ring of wild type (n=4) and IL-5-/- (n=4) mice following the application of vehicle (PBS) or denatonium. Data represent means ± SEM of 15-20 rings of 4 animals (B-C). One-way ANOVA followed by Bonferroni's multiple-comparison test (B-C).

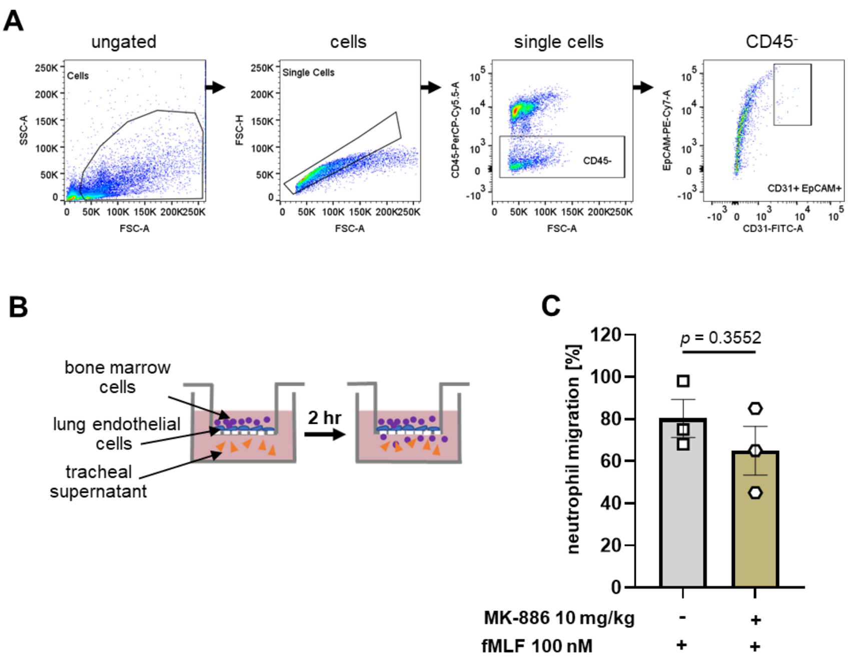


**Supplementary Figure 8: Transwell migration assay of bone marrow neutrophils.** (A) Gating strategy for flow cytometric identification of lung endothelial cells. Cellular debris was first excluded, and singlets were gated. Lung endothelial cells were identified as $CD45^-$ and $CD31^+$**.** (B) Demonstration of the transwell migration assay using FACS-sorted lung endothelial cells seeded on inserts. (C) Quantification of neutrophil migration in transwell assays toward N-formyl-l-methionyl-l-leucyl-l-phenylalanine (fMLF) (100 nM) (n=3). Data are means ± SEM of three independent experiments (C). Two-tailed unpaired Student's t-test (C).

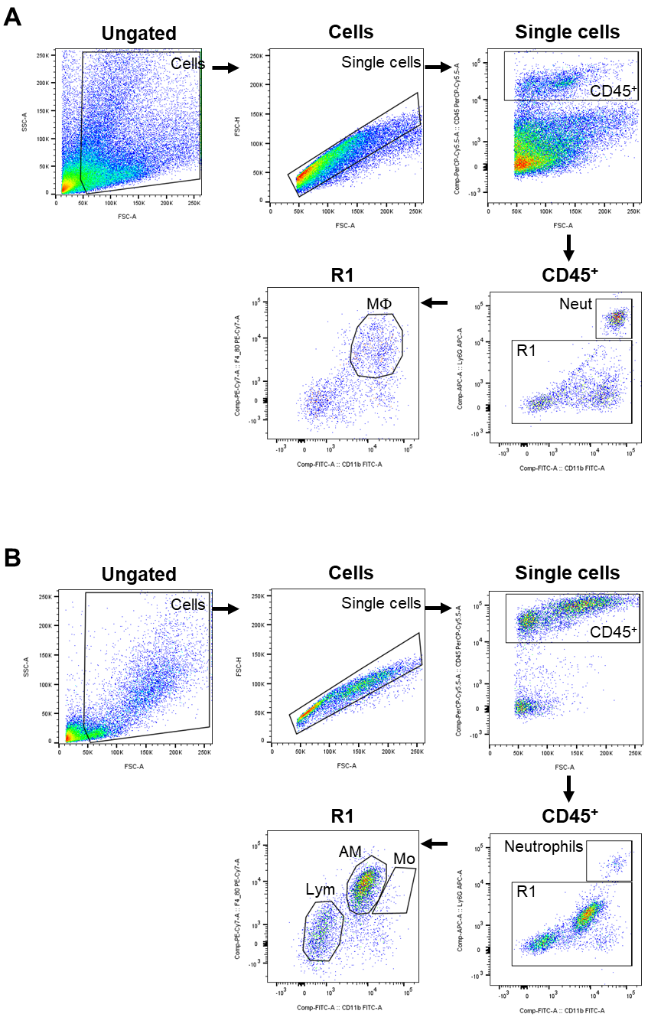


**Supplementary Figure 9: Gating strategies for tracheal (A) and BALF (B) immune cells identification by flow cytometry.** Cell debris, and doublets were excluded, and cells were identified as follows: neutrophil (Neut) ($CD45^+$ $CD11b^+$ $Ly6G^+$), macrophages/monocytes (MΦ/Mo) ($CD45^+$ $CD11b^+$ $Ly6G^-$ $F4/80^+$), alveolar macrophages (AM) ($CD45^+$, Ly6G-, $F4/80^+$ $CD11b^{low/-}$) and lymphocytes ($CD45^+$ $CD11b^-$ $Ly6G^-$ $F4/80^-$).

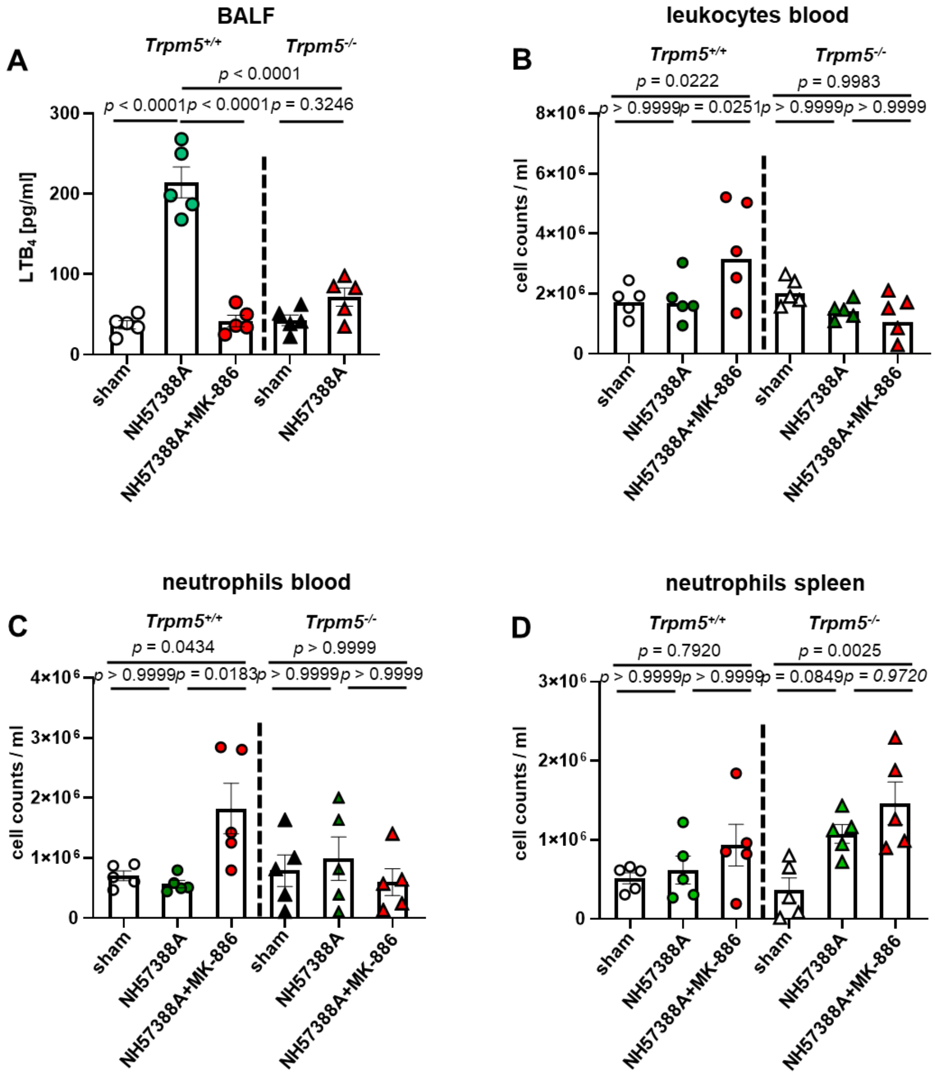


**Supplementary Figure 10: LTB4 mediates the tuft cell-induced recruitment of blood neutrophils and monocytes.** (A) ELISA measurement of LTB4 released in BALF retrieved from *Trpm5*$^{+/+}$ and *Trpm5*$^{-/-}$ mice, either sham-treated (n=5) or infected with the *P. aeruginosa* isolate NH57388A alone (n=5) or following the injection of MK-886 (10 mg/kg) (n=5). (B-D) FACS analysis of leucocytes (CD45+; B) and neutrophils (C) in the blood and neutrophils (D) in the spleen of infected animals (n=5). Data shown as means ± SEM of 4-5 independent experiments. One-way ANOVA followed by Bonferroni's multiple-comparison test.

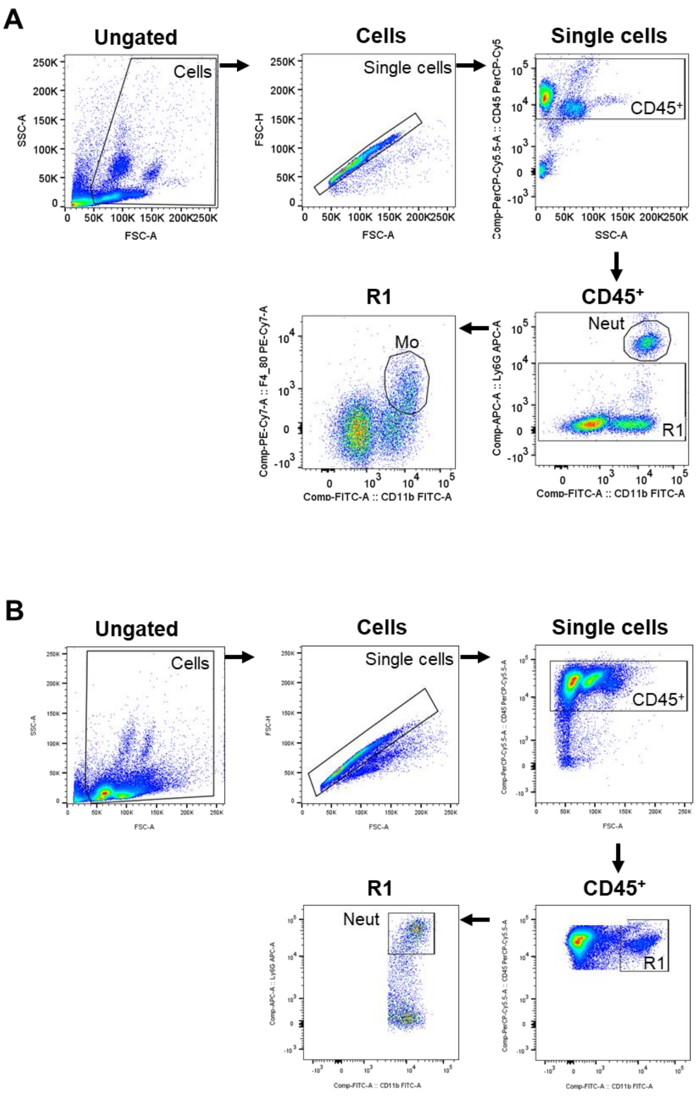


**Supplementary Figure 11: Gating strategies for blood (A) and spleen (B) immune cells identification by flow cytometry**. Following exclusion of cellular debris and doublets. Neutrophils (Neut) were identified as $CD45^+$ $CD11b^+$ $Ly6G^+$. Blood monocytes (Mo) were detected as $CD45^+$ $CD11b^+$ $Ly6G^-$ $F4/80^+$.

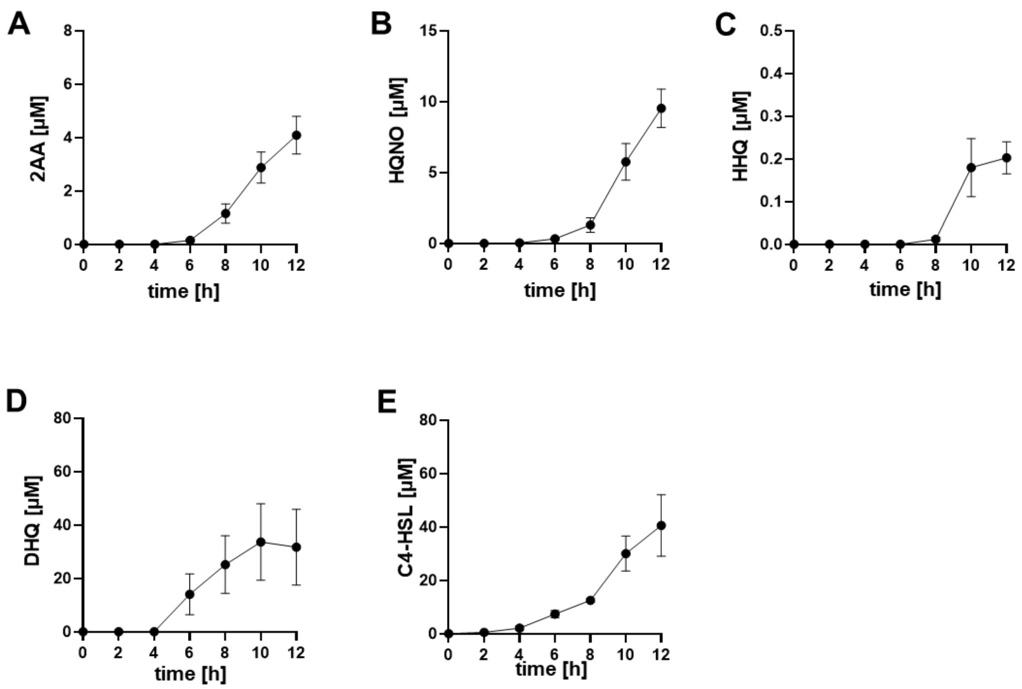


**Supplementary Figure 12: The *P. aeruginosa* isolate NH57388A releases quorum-sensing molecules (QSMs) during late phases of growth.** (A-E) Mass spectrometry measurement of QSMs (2-aminoacetophenone; 2AA, 2-heptylhydroxyquinoline N-oxide; HQNQ, 4-hydroxy-2-heptylquinoline; HHQ, 2,4-dihydroxyquinoline; DHQ, and N-Butanoyl-L-homoserine lactone; C4-HSL) released by the *P. aeuroginosa* strain NH57388A in LB medium at different time points of growth. Data represent means ± SEM of 3-5 independent experiments.

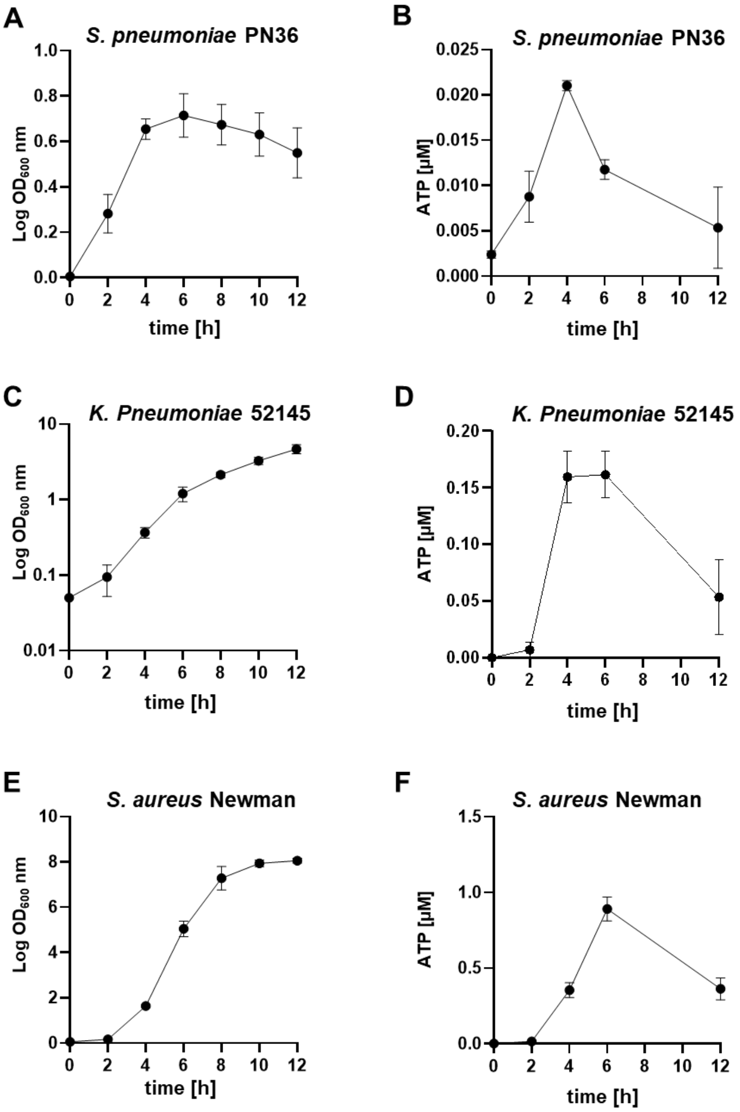


**Supplementary Figure 13: *Streptococcus pneumoniae* strain PN36, *Klebsiella pneumoniae* strain Kp52145, and *Staphylococcus aureus* strain Newman release ATP during growth.** (A) Growth curve of *Streptococcus pneumoniae* strain PN36 in THY medium (n=3). (B) Measurement of ATP released extracellularly in THY by *Streptococcus pneumoniae strain PN36* (n=3). (C) Growth curve of *Klebsiella pneumoniae* strain Kp52145 in LB medium (n=3). (D) Measurements of ATP released extracellularly by *Klebsiella pneumoniae* strain Kp52145 in LB. (E) Growth curve of S. aureus strain Newman in TSB (n=3). (F) Measurements of eATP released by the growing *S. aureus* strain Newman (n=3). Data represent means ± SEM of 3 independent experiments done in duplicates.

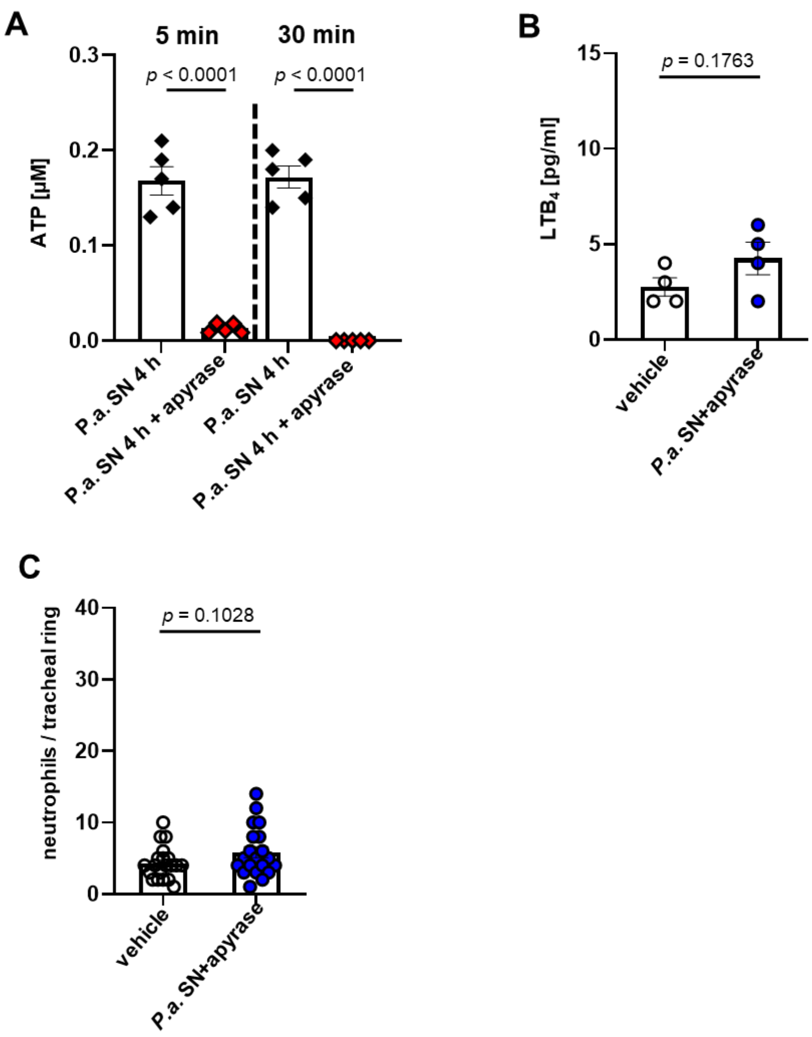


**Supplementary Figure 14: Bacterial eATP induces tuft cell-mediated recruitment of neutrophils via LTB4.** (A) Measurement of extracellular ATP in bacterial supernatants from *P. aeruginosa* incubated with apyrase (5 U/ml) for 5 and 30 min. (B) ELISA measurement of LTB4 released from tracheal tufts cells in response to *P. aeruginosa* supernatants collected at 4 hrs after growth and mixed with apyrase (5 U/ml) (n=4). (C) Quantification of neutrophils, per tracheal ring, 30 min after the intratracheal administration of *P. aeruginosa* supernatants either alone or mixed with apyrase (5 U/ml) (n=4). Data represent means ± SEM of 5 biological replicates measured in duplicates (A), 4 mice (B), and 20 tracheal rings obtained from 4 mice (C). One-way ANOVA followed by Bonferroni's multiple-comparison test (A-C).

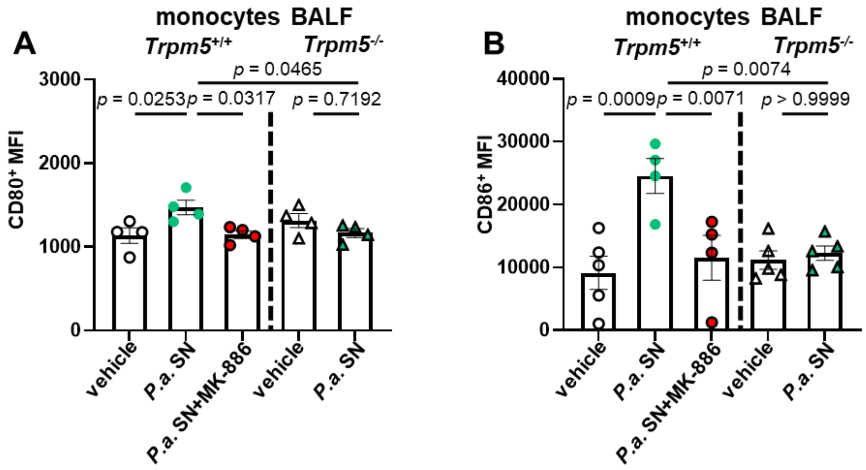


**Supplementary Figure 15: Tuft cell-derived LTs in response to bacterial supernatants drive monocyte activation.** (A-B) Quantification of the mean fluorescence intensity (MFI) of CD80 and CD86 in BALF of *Trpm5*$^{+/+}$ (n=4-5) and *Trpm5*$^{-/-}$ mice (n=4-5) 30 min after intratracheal treatment with vehicle (medium) or NH57388 supernatants (*P.a.* SN, 4 h). Data represent mean ± SEM of 4-5 mice (A-E). One-way ANOVA followed by Bonferroni's multiple-comparison test (A-E).

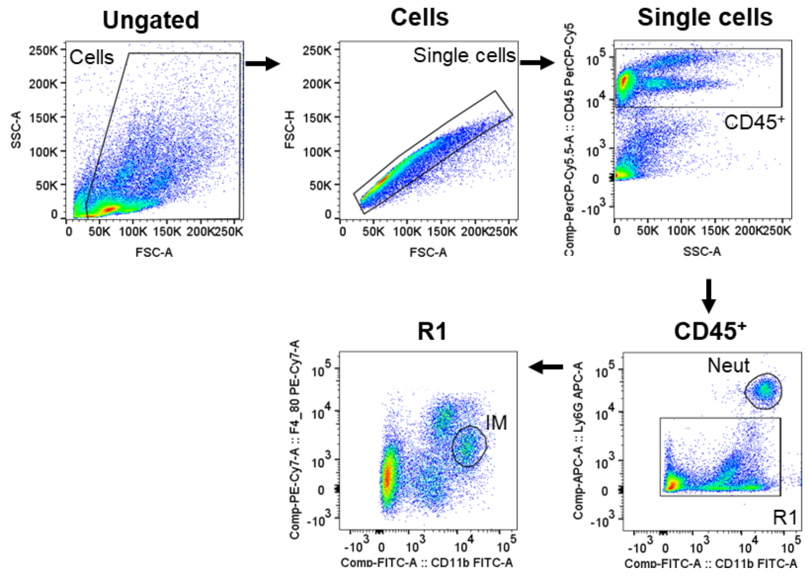


**Supplementary Figure 16: Gating strategy for lung interstitial macrophages FACS sorting.** After excluding the cell debris and doublets, lung interstitial macrophages (IM) were identified as $CD45^{+}$ $CD11b^{+}$ Ly6G- F4/$80^{+}$.

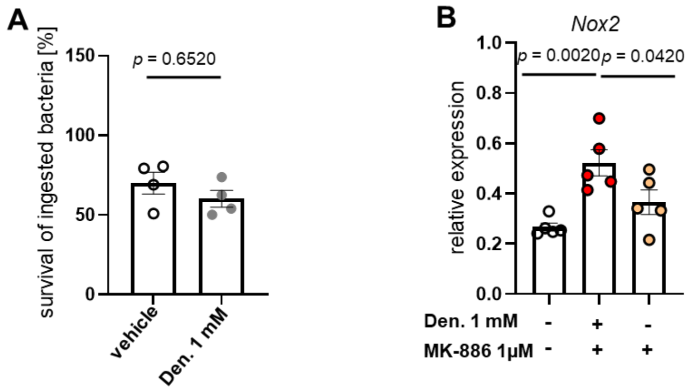


**Supplementary Figure 17: Tuft cell activation contributes to the production of hydrogen peroxide ($H_2O_2$) by macrophages.** (A) Bacterial killing capacity of lung interstitial macrophages exposed to 1mM denatonium before infection with *P. aeuroginosa* NH57388A (n=4) (B) qRT-PCR of *Nox2* expressed by lung interstitial macrophages upon exposure to tracheal supernatants from tracheae stimulated with denatonium. When required, the synthesis of LTs was inhibited by preincubating the trachea with MK-886 (n=5). Data shown as mean ± SEM of 4-5 independent experiments. Two-tailed unpaired Student's t-test (A) or One-way ANOVA followed by Bonferroni's multiple-comparison test (B).
